# A computational study about the mechanism of action of metformin on hepatic gluconeogenesis, focused on its ability to create stable pseudo-aromatic copper complexes

**DOI:** 10.1101/130211

**Authors:** Fabio Rebecchi

## Abstract

Metformin is the best therapeutic choice for treating type 2 Diabetes.

Despite this, and the fact it has been prescribed worldwide for decades, its mechanism of gluconeogenesis inhibition is still unknown.

In the following work a novel mechanism of inhibition is suggested: that metformin performs its action on the target enzyme not as a pure molecule but, after sequestering endogenous cellular copper, as a copper complex.

This result was obtained using chemoinformatics methods including homology modeling for the creation of the target enzyme’s tridimensional virtual structure, molecular docking for both the determination of the movement of the prosthetic group inside its cavity and for the identification of the best ligand poses for the metformin copper complexes, and eventually pharmacophore modeling and virtual screening to find alternative virtual leads that could achieve similar effects.

The simulations show the complex binding as a non competitive inhibitor to the large exit of the mitochondrial glycerophosphate dehydrogenase enzyme’s FAD cavity, preventing FAD movement inside the cavity and/or quinone interaction and therefore its electron transfer function.

The proposed mechanism seems to be successful at explaining a wide range of existing experimental results, both regarding measurements of metformin non-competitive inhibition of GPD2 and the role of copper and pH in its action.

The virtual screening outcome of at least two similarly active purchasable molecule hints to an easy way to experimentally test the proposed mechanisms.

In fact, the virtual leads are very similar to the copper complex but quite different from metformin alone, and a laboratory confirmation of their activity should plausibly imply that metformin acts in synergy with copper, giving us the ability to design new antidiabetic drugs in a novel and more rational fashion, with significant savings in research costs and efforts.

## Section 1 Introduction

### 1.1 Metformin: a brief story

Metformin is a molecule that belongs to the biguanidine class of drugs (Figure 1.1).

**Figure 1.1.**
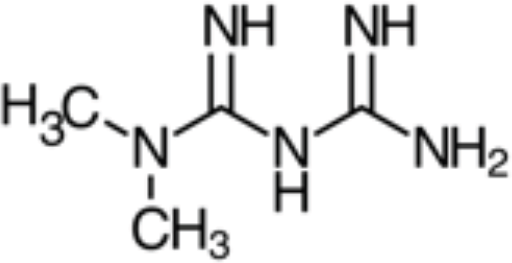
Structural formula of metformin.

Its hypoglycemic effect in animals has been well known since 1929, and the same effect on humans was discovered in 1962. (Gottlieb & Auld, 1962).

It is the first therapeutic resource for Type 2 Diabetes Mellitus, being able to inhibit hepatic gluconeogenesis, and preventing hyperglycemia, without causing increase in insulin secretion, hypoglycemia or weight gain. (Bennet et al., 2010)

It has been prescribed for over forty years now (Rojas, Chávez-castillo & Torres, 2014).

Furthermore, metformin showed the ability to decrease the incidence of several kind of cancers in diabetic patients who underwent metformin therapy (Karabulut-Bulan, 2016; Rojas, Chávez-castillo & Torres, 2014), and it recently attracted the attention of anti-ageing researchers because it has been noted to extend lifespan in nematodes and mice (Onken & Driscoll, 2010).

### 1.2 Metformin and gluoconeogenesis: a review

In spite of the long and widespread use of metformin, its mechanism of action is unknown. Through the years, many investigations addressed the problem, often with contradictory results. The earliest studies focused on respiratory Complex I, and (Owen & Halestrap, 2000) found that millimolar concentrations of metformin mildly inhibit respiratory chain’s Complex I in isolated mitochondria and this effect seemed to be directly linked to the gluconeogenesis inhibition. Then, researchers shifted their attention to the LKB1/AMPK pathway, with (Zhou et al, 2001) reporting that metformin needs AMPK to exert its inhibitory effect on gluconeogenesis, taking place through AMPK inhibition of SREBP-1 gene expression, and hypothesized that metformin directly activates AMPK. On the road traced by these results a subsequent study (Shaw et al, 2005), pointed out that metformin needs LBK1, the AMPK upstream activator, to decrease blood glucose level.

However, (Savage et al, 2006) and (Fullerton et al, 2013) showed that the inhibition of gluconeogenic transcription could be achieved by a parallel mechanism also, i.e. it could be due to the sensitization of transcription to insulin where metformin activity depends on ACC and induces an AMPK/ACC mediated decrease of hepatic lipid content.

Furthermore, the AMPK signalling inhibition hypothesis is contradicted by the results of (Foretz et al, 2010), that found that metformin depended solely on the ability to decrease hepatic energy charge to exert its anti-hyperglicemic action, performed even in mice lacking AMPK or LKB1, and by the results of (Miller et al, 2013), showing that metformin increases cAMP concentration and cAMP is capable of inhibiting adenylyl cyclase preventing its activation by glucagon and therefore gluconeogenesis as well.

But the reduction of hepatic energy charge’s hypothesis has its opponent too.

(Hawley, Olsen & Hardie, 2002) in fact, showed not only that metformin does not affect regulation of AMPK by upstream kinases and phosphatases, but also that it activates AMPK without altering the AMP/ADP and the ADP/ATP ratios.

In this intricate and contradictory scenario, a recent study (Madiraju et al, 2014) took into consideration all the previous results listed above and suggested, backed by original and exhaustive experimental data, that metformin actually reduces gluconeogenesis by non-competitive inhibition of the mitochondrial glycerophosphate dehydrogenase (GPD2).

Supplementary insight on the issue is given by four other studies that target the interaction of biguanides with copper and the metal role in their action.

(Repiščák et al., 2014) shows that, among biguanides, metformin is not only the best at inhibiting gluconeogenesis, but it’s the most effective at forming stable pseudo-aromatic copper complexes, suggesting a correlation between the two facts. Moreover, the complex stability increases with deprotonation, an event that is more likely to happen in the high pH environment of mitochondria. (Logie et al., 2012) found that glucose production in liver cells is no longer decreased by metformin when cellular copper is sequestered, suggesting that copper is essential to metformin action. (Annapurna & Rao, 2007) found that, in alloxan induced diabetic rabbits, Cu^2+^ and Ni^2+^ metal complexes of metformin showed a significant increase in hypoglicemic activity when compared to the pure drug. This pattern seems common in biguanides, since the antiproteolytic action of metformin and phenformin shows a significant increase for the metal complexes when compared to pure drugs (Sweeney et al, 2003).

### 1.3 Project objectives and scope

(Madiraju et al., 2014) proposes a mechanism of action based on the hypothesis that FAD, the GPD2’s prosthetic group, needs to perform an oscillatory sliding-in/sliding-out movement in order to first oxidise glycerol-3-phosphate and then transfer the electrons through the transport chain, and that metformin blocks this movement by docking with the smaller of the cavity’s two entrances.

But a pilot simulation carried out as part of this work, performed using molecular docking software on 2RGH, the bacterial alpha-glycerophosphate oxidase that is a template for GPD2, showed three weaknesses in this hypothesis.

The first was that the free binding energy of metformin docked in the cavity’s small entrance is too little to account for the inhibition constant (KI) found in (Madiraju et al., 2014).

The second was that metformin seems to bind to this target only when glycerol-3-phosphate, its substrate, is absent, therefore lacking one of the required characteristics of a non-competitive inhibitor.

The third was that this mechanism isn’t able to explain why the absence of copper halts the inhibitory action.

The same simulation showed that the weaknesses above are overcome when a square planar copper- complex of metformin (Figure 1.2) is used as a ligand instead, this time with a preferential docking area in the cavity’s large entrance.

**Figure 1.2.**
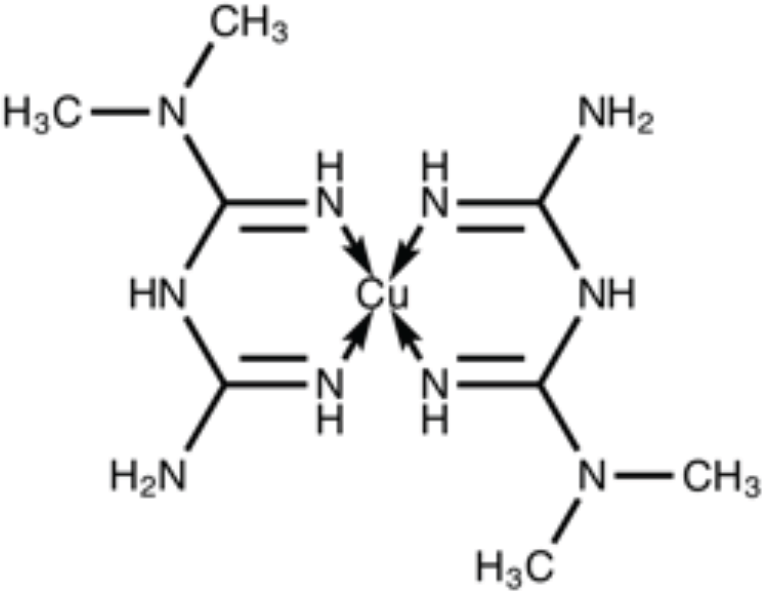
Structural formula of a metformin Cu^2+^copper complex.

This preliminary result seemed to suggest that metformin performs its inhibitory action by collecting endogenous cellular copper and by docking on GPD2 as a copper-complex, and this mechanism could explain not only the inhibitory effects measured in (Madiraju et al., 2014) but account at the same time both for the role of copper in the process as highlighted in (Logie et al., 2012) and for the increased potency of the metal complex shown in (Annapurna & Rao, 2007).

This prompted to carry out a systematic series of molecular docking simulations aimed at understanding in depth the multiple interactions between GPD2, FAD, glycerol-3-phosphate, metformin alone, monometformin copper complex, dimetformin copper complex and coenzyme Q10, the final electron acceptor. Simulation runs were performed with phenformin too, a similar antihyperglycemic biguanide.

Objectives of these simulations were to identify the most favourable FAD positions, both alone and bond with glycerol-3-phosphate, and the most favourable poses for metformin and its complex to achieve inhibition by blocking the FAD’s electron transport function.

The best docking poses, achieved by the copper complex, were then exploited to perform a virtual screening on databases of commercial molecules with the aim of identifying lead compounds able to exert a similar action.

## Section 2: Methods

### 2.1 Protein structures retrieval

PDB structure of 2RGH was retrieved from Protein Data Bank (Berman et al., 2000).

GPD2 amino acid sequence in FASTA format was retrieved from UniProt (The UniProt Consortium, 2014).

### 2.2 Ligand structures retrieval

PDB structures of FAD, G3P and co-enzyme Q10 were retrieved from Protein Data Bank (Berman et al., 2000).

PDB structures of metformin and phenformin were retrieved from Drug Bank (Wishart et al., 2006). The lead candidates structures were obtained in MOL2 file format from Zinc (Irwin et al., 2012) and from ChemSpider (Royal Society of Chemistry, 2016), and subsequently converted into PDB by Open Babel (O’Boyle et al., 2011), and eventually refined in Discovery Studio (Dassault Systèmes, 2016).

### 2.3 Protein structures modeling

Since GPD2’s crystallographic structure was unavailable, the GPD2 FAD cavity was modeled by substituting the 2RGH cavity residues with the corresponding residues of the GPD2 amino acid sequence.

First, in order to achieve the original protein structure, as it was prior to the processing needed for the crystallography procedure, the seleno-methionine residues of the 2RGH structure were substituted with methionine residues using the Macromolecules/replace function of Discovery Studio (Dassault Systèmes, 2016).

The residues forming the FAD cavity in 2RGH were identified using Receptor-Ligand Interaction Explorer of Discovery Studio (Dassault Systèmes, 2016).

The other cavity residues, corresponding to the small and the large cavity entrance, were manually identified in Autodock Tools (Morris et al., 2009).

The correspondence between the GPD2 and the 2RGH residues was determined using the target-template alignment function present on I-Tasser (Yang et al., 2015) that returned 2RGH as one of the top threading templates for the GPD2’s FASTA sequence in input (Figure 2.1).

**Figure 2.1.**
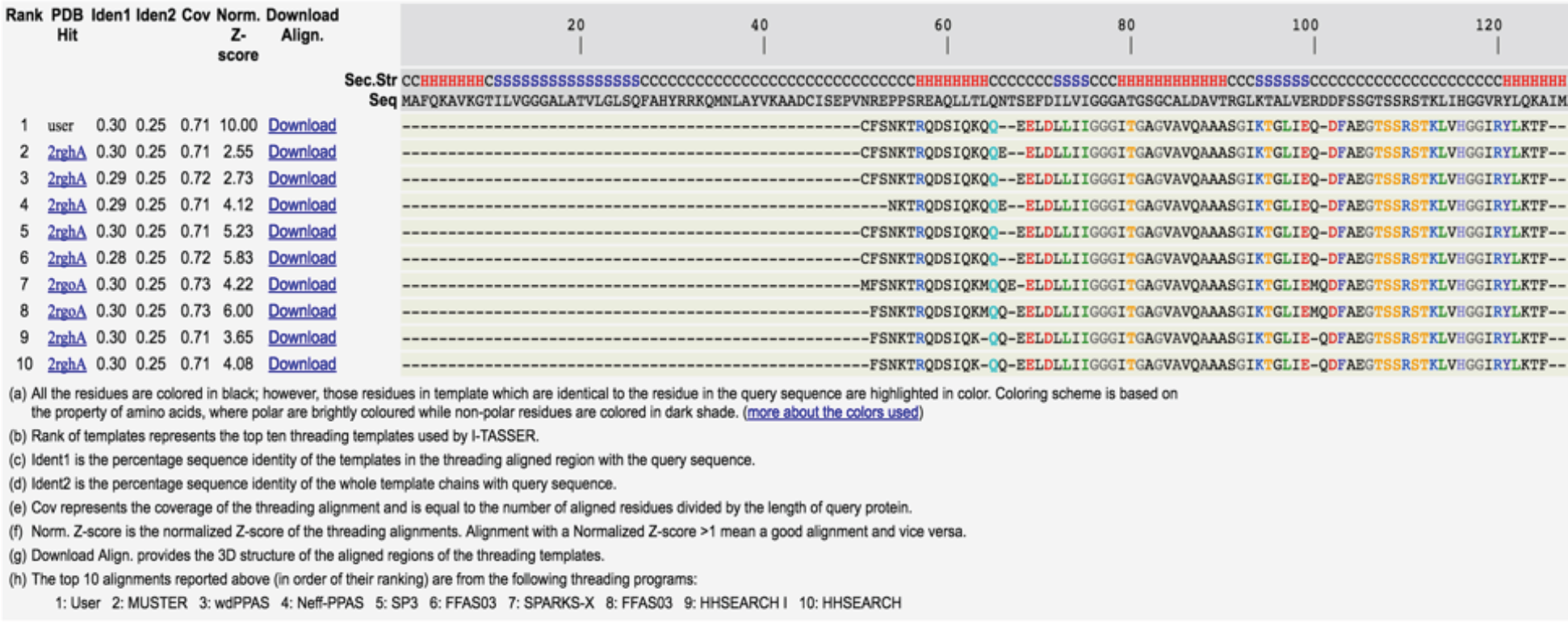
The correspondence between the GPD2 residues and the residues of 2RGH, the best threading template in I-Tasser.

The substitution of the 2RGH cavity residues with the corresponding GPD2 residues was performed with Discovery Studio (Dassault Systèmes, 2016) using the Macromolecules/replace function.

### 2.4 Ligand structures modeling

The dimetformin (Figure 2.2) square planar copper complex was created in Discovery Studio (Dassault Systèmes, 2016) using the bond distances reported in (Repiščák et al., 2014).

**Figure 2.2.**
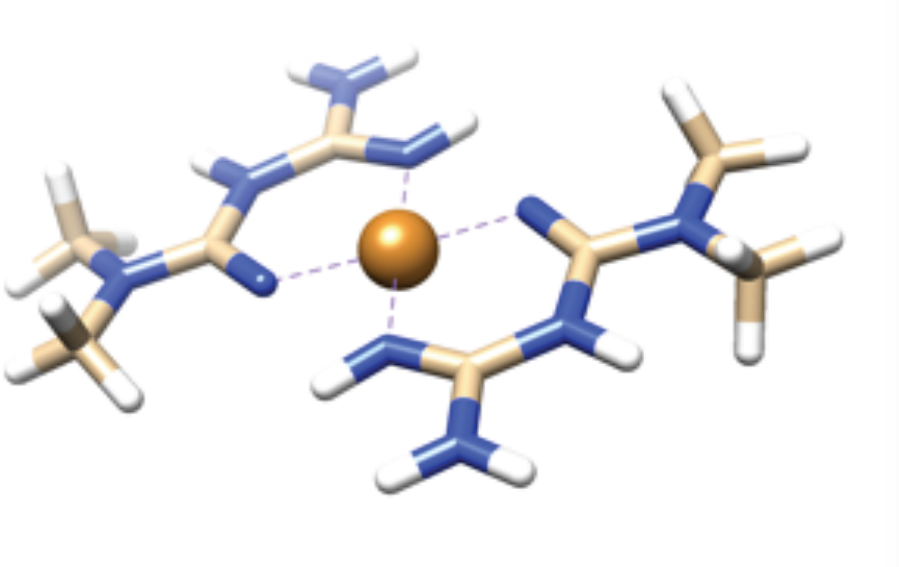
A ligand modeling example: the dimetformin neutral copper complex visualised in Chimera.

The protonation states of residues and ligands at various pHs (Figure 2.3). were determined by MarvinBeans (ChemAxon, 2016).

**Figure 2.3.**
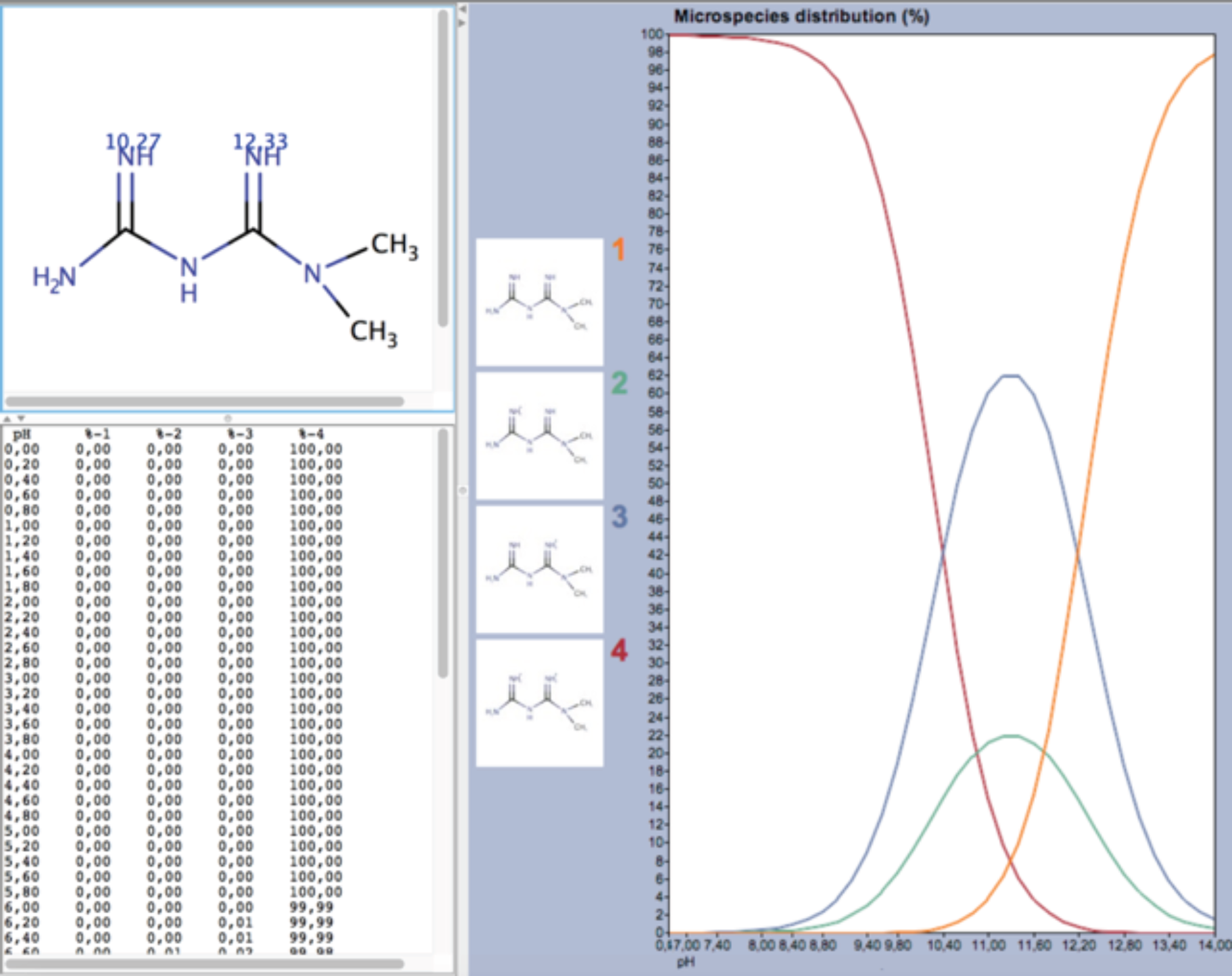
An example of speciation curves obtained by MarvinBeans, in this case for metformin; species 1 is non-protonated, species 2 and 3 are mono-protonated, species 4 is di-protonated.

### 2.5 Molecular docking

A first series of molecular dockings were performed for FAD into the GPD2’s cavity, in order to assess the plausibility and the extent of a FAD movement inside it.

Then glyceraldehyde 3-phosphate (G3P) was docked with various positions of FAD to assess the most suited FAD positions for docking G3P, and coenzyme Q10 was docked as well to understand its target site for electrons retrieval.

Eventually, the following ligands were docked to the protein with FAD in different positions, to assess their ability to act as non competitive inhibitors: pure metformin, pure phenformin, dimetformin copper complex,, dimetformin copper complex, monophenformin copper complex, and the two alternative lead candidates described in the Results section.

Molecular docking of ligands on the GPD2 cavity was performed using Autodock 4 (Morris et al., 2009) and Autodock Vina (Trott & Olson, 2010).

On Autodock 4, a parameter file with atomic data for copper was added, since copper wasn’t supported in the standard version.

PDBQT format files for ligands and for the protein structure, necessary for Autodock input, were prepared by adding hydrogens, computing Gasteiger charges (Gasteiger & Marsili, 1980) and merging non polar hydrogens on Autodock Tools (Morris et al., 2009). Where necessary, bonds to be considered flexible in ligands and residues were set up on PDBQT files using Autodock Tools (Morris et al., 2009). The total number of flexible bonds in a simulation ranged from zero to 25.

The coordinates Grid Boxes (Figure 2.4), necessary to delimit the simulation docking area, were prepared using Autodock Tools (Morris et al., 2009). Grids’ volume was between 1000 and 2700 cubic angstroms.

**Figure 2.4.**
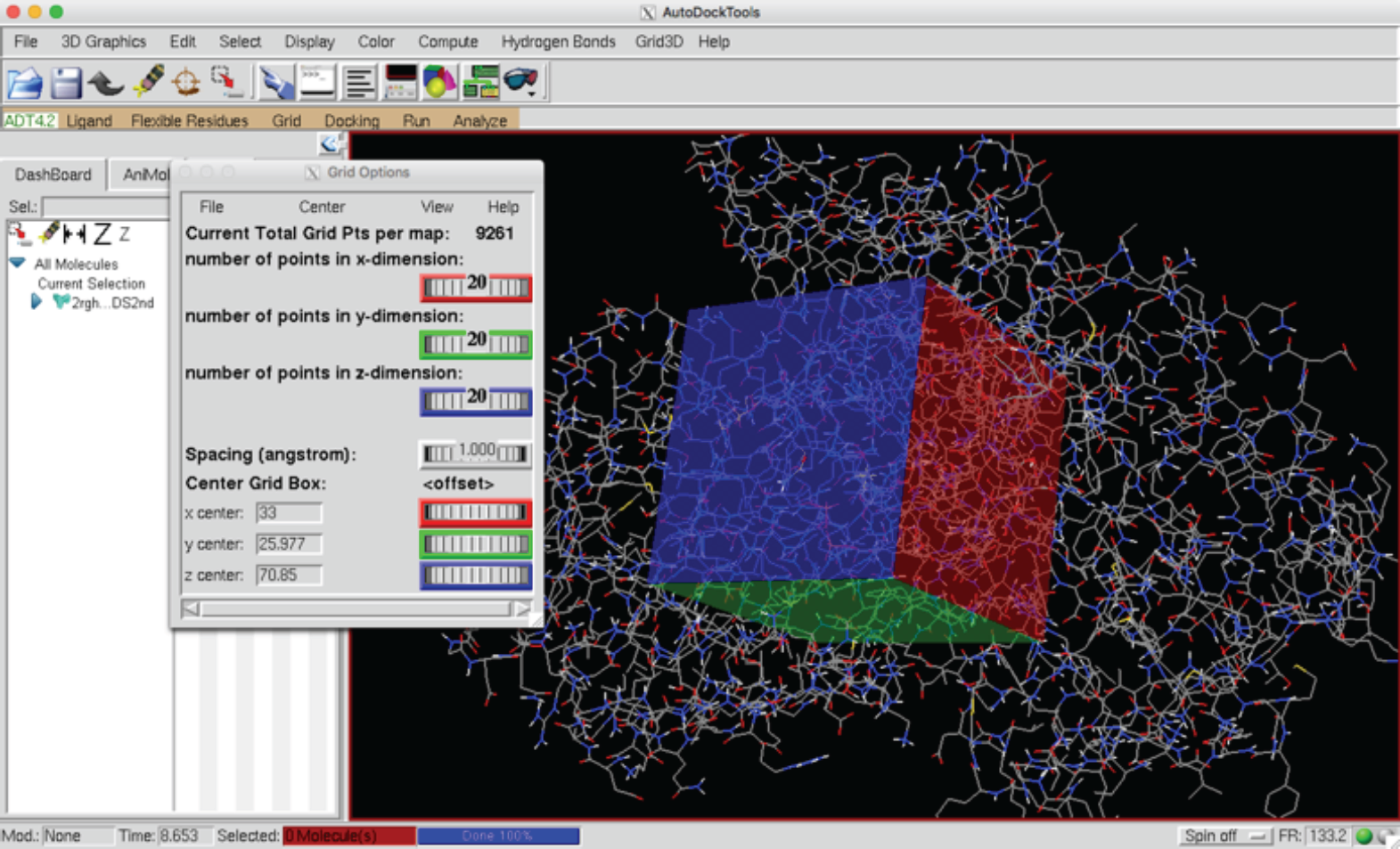
The set up of a coordinates grid (Grid Box) on Autodock Tools to delimit the docking area.

The searches were set up with a higher level of accuracy than standard, at the expense of calculation speed, with 25 millions of maximum evaluations set on Autodock 4 (Morris et al., 2009) and the exhaustiveness parameter set on 25 on Autodock Vina. (Figure 2.5)

**Figure 2.5.**
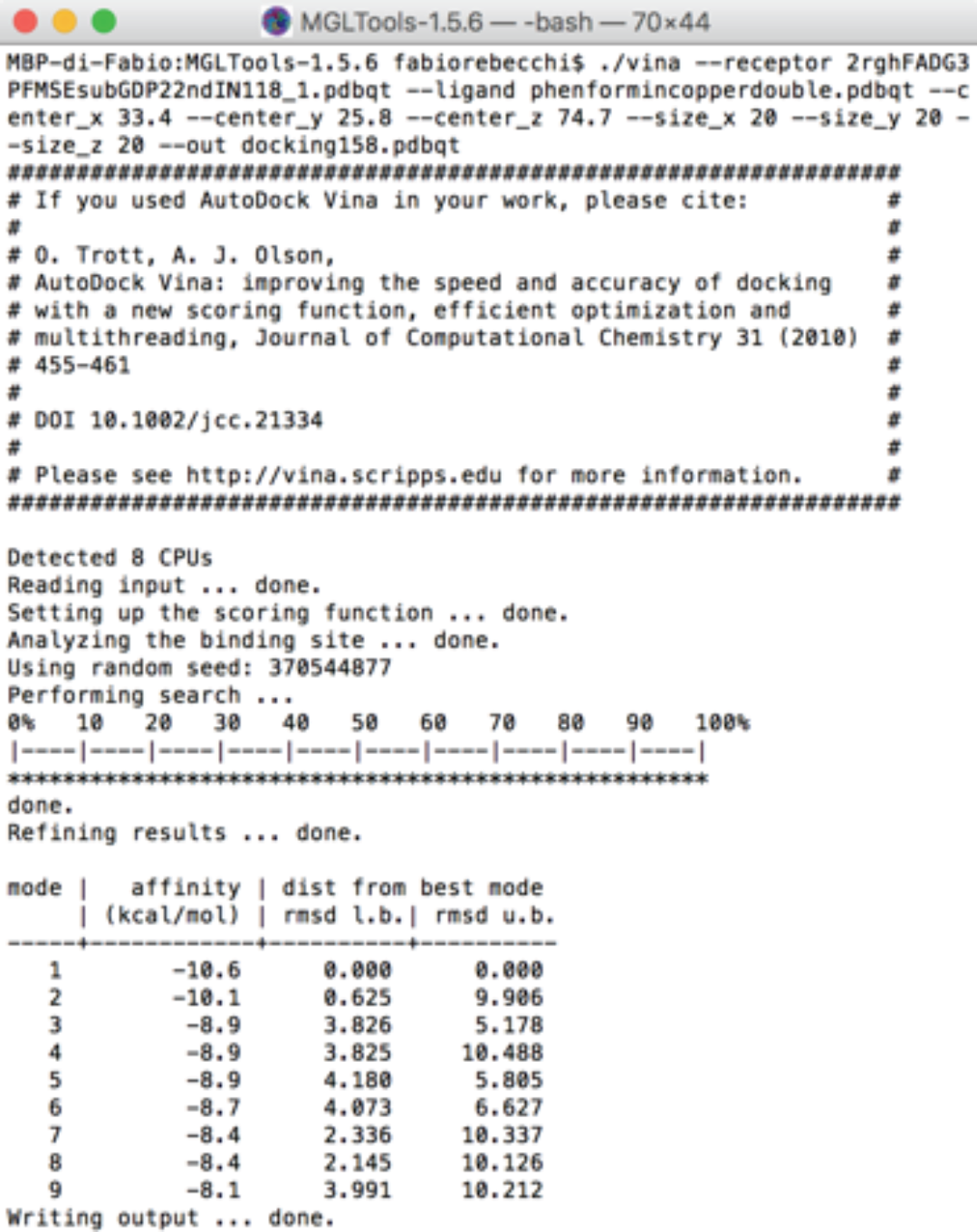
An example of input/output screen for Autodock Vina inside the command line shell.

The number of poses to be returned was set on 50 inside an energy range of maximum 50 kcal/mol. The inhibition constant Ki was calculated as nanomoles applying the following formula: 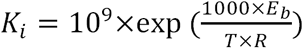,

where E_b_ is the binding energy expressed in kcal/mol, T is the temperature (298K) expressed in Kelvins and R is the gas constant approximated as 1.987 cal K^-1^ mol^-1^.

### 2.6 Analysis of ligand interaction and binding cavities

A preliminary control of results was performed using Autodock Tools (Morris et al., 2009) and Chimera (Pettersen et al., 2004).

The analysis of the docked poses and of the interaction of the ligands with the cavity residues was performed using the Receptor-Ligand Interaction Explorer of Discovery Studio (Dassault Systèmes, 2016).

### 2.7 Pharmacophore and virtual screening

A pharmacophore descriptor based on the dimetformin copper complex was created using the manual pharmacophore creation feature (Figure 2.6) of ZincPharmer (Koes & Camacho, 2012).

**Figure 2.6.**
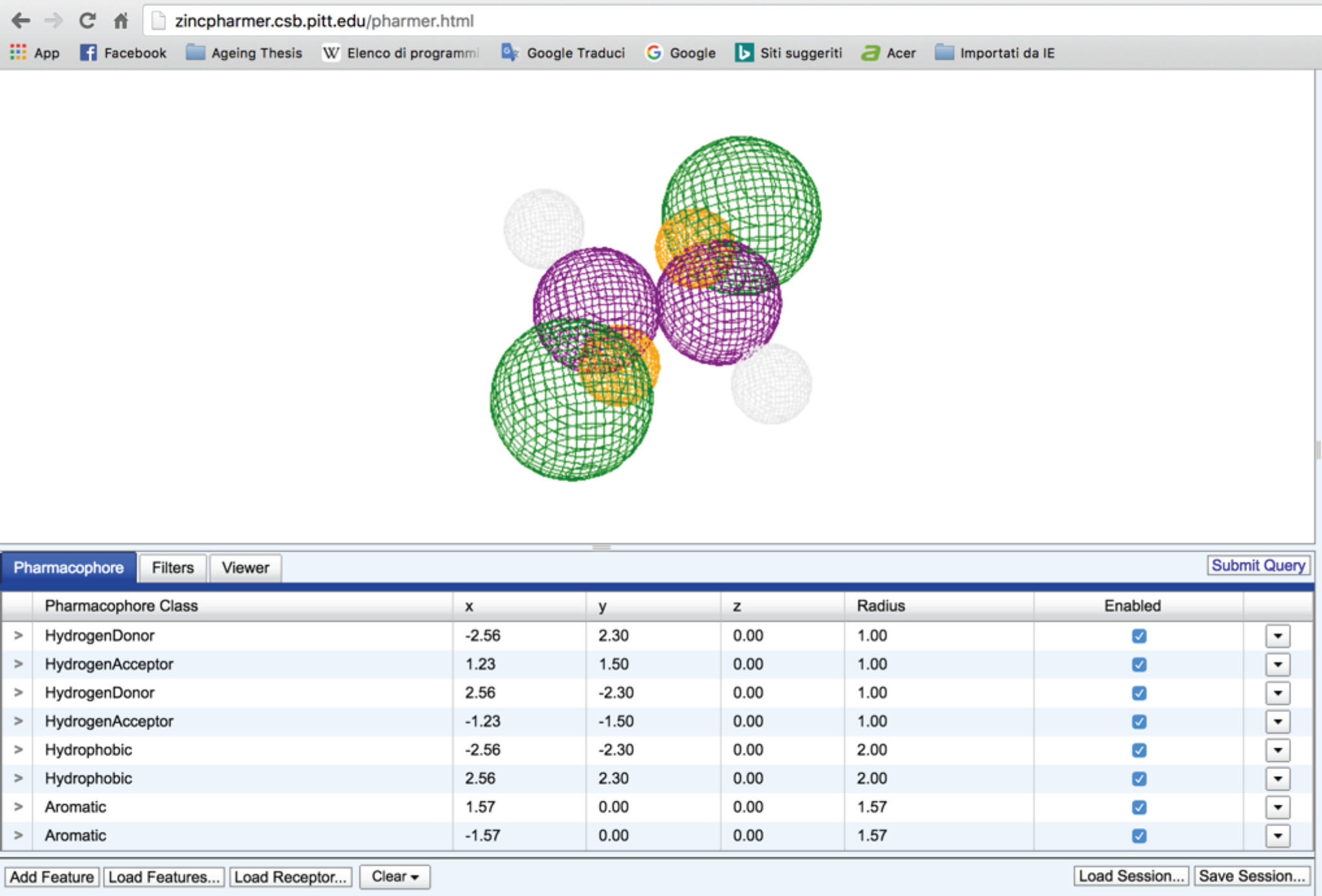
Pharmacophore modeling interface of the ZincPharmer website.

The coordinates and radius of each functional group were derived from distance measurements taken in Discovery Studio (Dassault Systèmes, 2016) on the copper complex PDB model.

A virtual screening was performed on the “Purchasable Compounds” of Zinc (Irwin et al., 2012). The parameters used for the search were the following:

- Molecular Weight: less than 250
- Max hits 100: 1
- Max hits per conf: 1
- Max hits per mol: 1

Two new compounds were designed from scratch too, using MarvinSketch (ChemAxon, 2016), following simple similarity considerations with the monometformin and the dimetformin copper complexes. The compounds are described in the Results section.

In order to obtain other lead candidates, a similarity search on these new compounds was performed on ChemSpider (Royal Society of Chemistry, 2016),

### 2.8 Hardware used

The following hardware was employed for Molecular Docking:

A MacBook Pro with an 8-core Intel i7 2.8 GHZ CPU and 16 GB 1600 MHz DDR3 RAM. The following hardware was employed for all other activities:

An Alienware PC with an Intel Core i7-6820HK (Quad-Core, 8MB Cache, 4.1GHz) CPU and 32 GB RAM.

## Section 3 Results

### 3.1 Homology modeling

The analysis of the FAD binding cavity of 2RGH showed it is made up of the 66 residues reported in Table 3.1.

**Table 3.1.** List of the residues forming the original 2RGH cavity.

| List of 2RGH's residues |
| --- |
| GLY25, GLY26, GLY27, ILE28, THR29, ILE47, GLU48, MET49, GLN50, GLY55, THR56, SER57, ARG59, SER60, THR61, LEU63, VAL64, HIS65, ILE68, ARG69, TYR70, MET78, VAL79, MET195, LYS196, ALA197, VAL198, THR231, SER232, GLY233, PRO234, TRP235, VAL236, ASP237, LYS238, VAL239, ARG240, ASN241, LEU242, THR255, LYS256, GLY257, ILE258, HIS259, TYR274, PHE275, ASP276, THR277, GLY282, ARG283, MET284, VAL285, THR298, ASP300, THR301, GLY344, LEU345, ARG346, PRO347, LEU348, VAL349, GLY428, LYS429, ILE430, THR431 |

This result includes the edges of both the small and the large entrance.

Confronting the above residues with the corresponding ones in the GPD2 sequence gave the result highlighted in Figure 3.1, with 24 residues not matching one another, corresponding to the 36% of the total.

**Figure 3.1.**
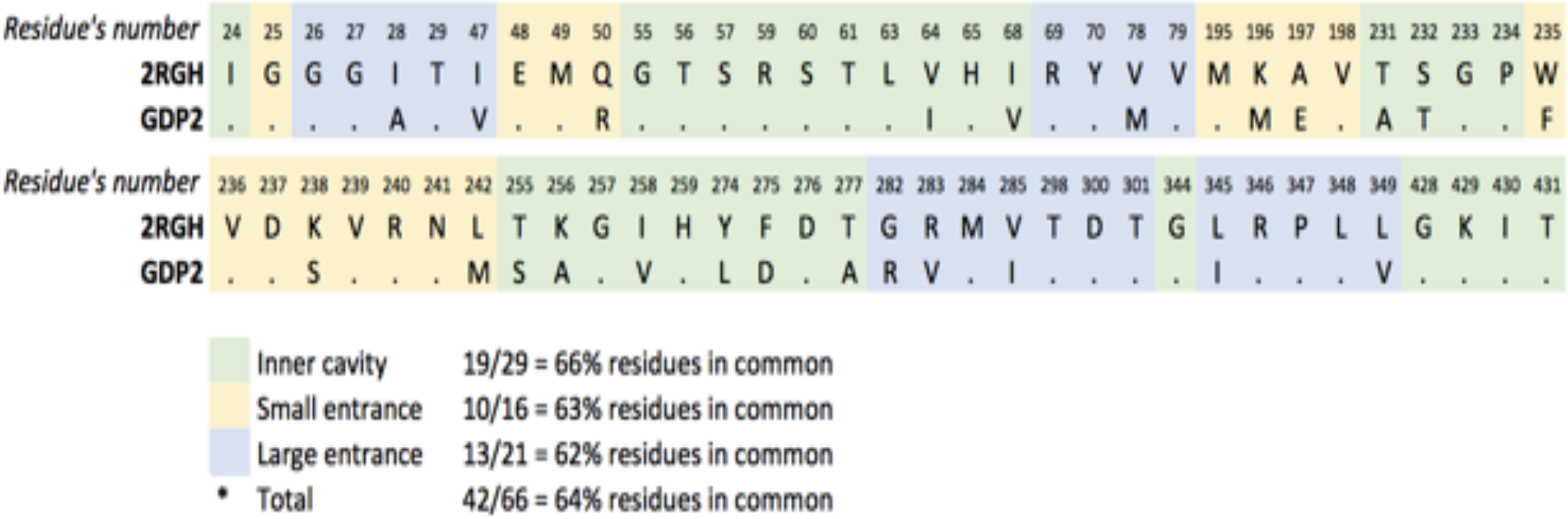
Cavity’s residues comparision between 2RGH and GPD2.

After substituting into the 2RGH tridimensional structure the non matching residues from GPD2, the binding cavity 66 residues became the ones reported in Table 3.2.

**Table 3.1.**
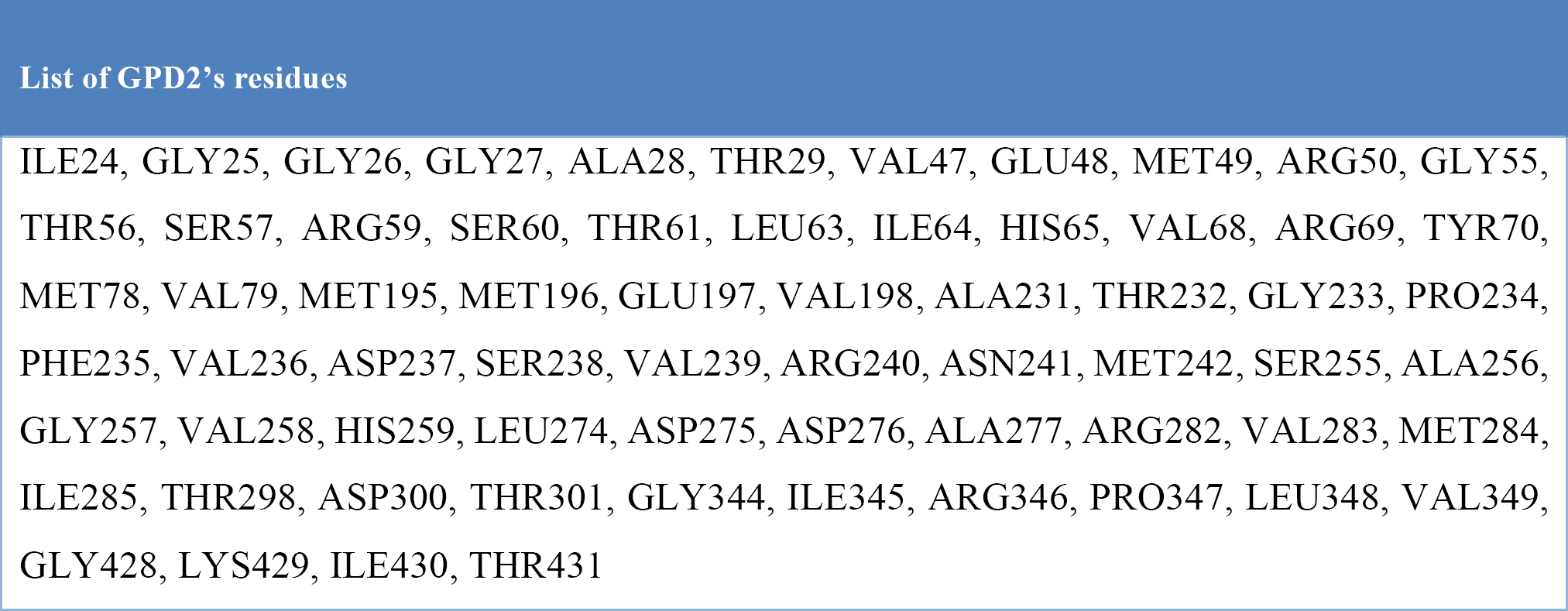
List of the residues forming the modeled GPD2 cavity.

| List of GPD2's residues |
| --- |
| ILE24, GLY25, GLY26, GLY27, ALA28, THR29, VAL47, GLU48, MET49, ARG50, GLY55, THR56, SER57, ARG59, SER60, THR61, LEU63, ILE64, HIS65, VAL68, ARG69, TYR70, MET78, VAL79, MET195, MET196, GLU197, VAL198, ALA231, THR232, GLY233, PRO234, PHE235, VAL236, ASP237, SER238, VAL239, ARG240, ASN241, MET242, SER255, ALA256, GLY257, VAL258, HIS259, LEU274, ASP275, ASP276, ALA277, ARG282, VAL283, MET284, ILE285, THR298, ASP300, THR301, GLY344, ILE345, ARG346, PRO347, LEU348, VAL349, GLY428, LYS429, ILE430, THR431 |

Figure 3.2 shows the cavity and the position of the small and of the large entrance in the newly modeled cavity.

**Figure 3.2.**
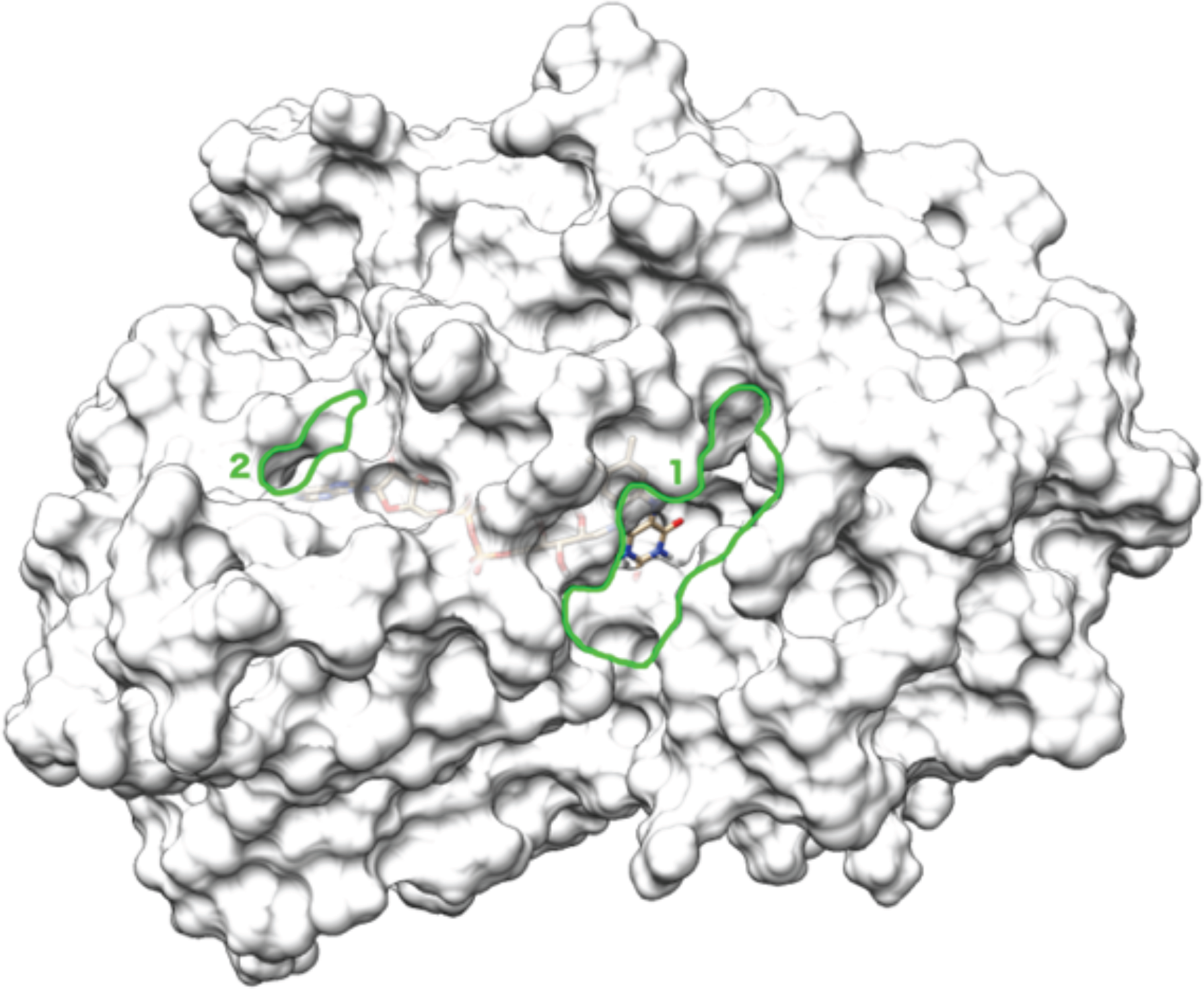
The newly modeled cavity structure with docked FAD, the edges of the small (2) and large (1) entrances are highlighted in green. The white hydrophobicity surface has 50% transparency, showing the FAD body inside.

### 3.2 Molecular docking

#### 3.2.1 FAD sliding movement

The redocking of FAD (Figure 3.3) in the new cavity gave a best pose with a RMSD of atomic positions less than 0.2 angstroms from the original 2RGH crystal structure pose (Colussi et al., 2008), as shown in Figure 3.4, confirming a good reliability of the docking software. The estimated free binding energy of the original FAD from 2RGH crystallography was 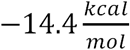, while the energy of the best pose for the FAD docked in the new cavity was 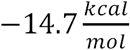.

**Figure 3.3.**
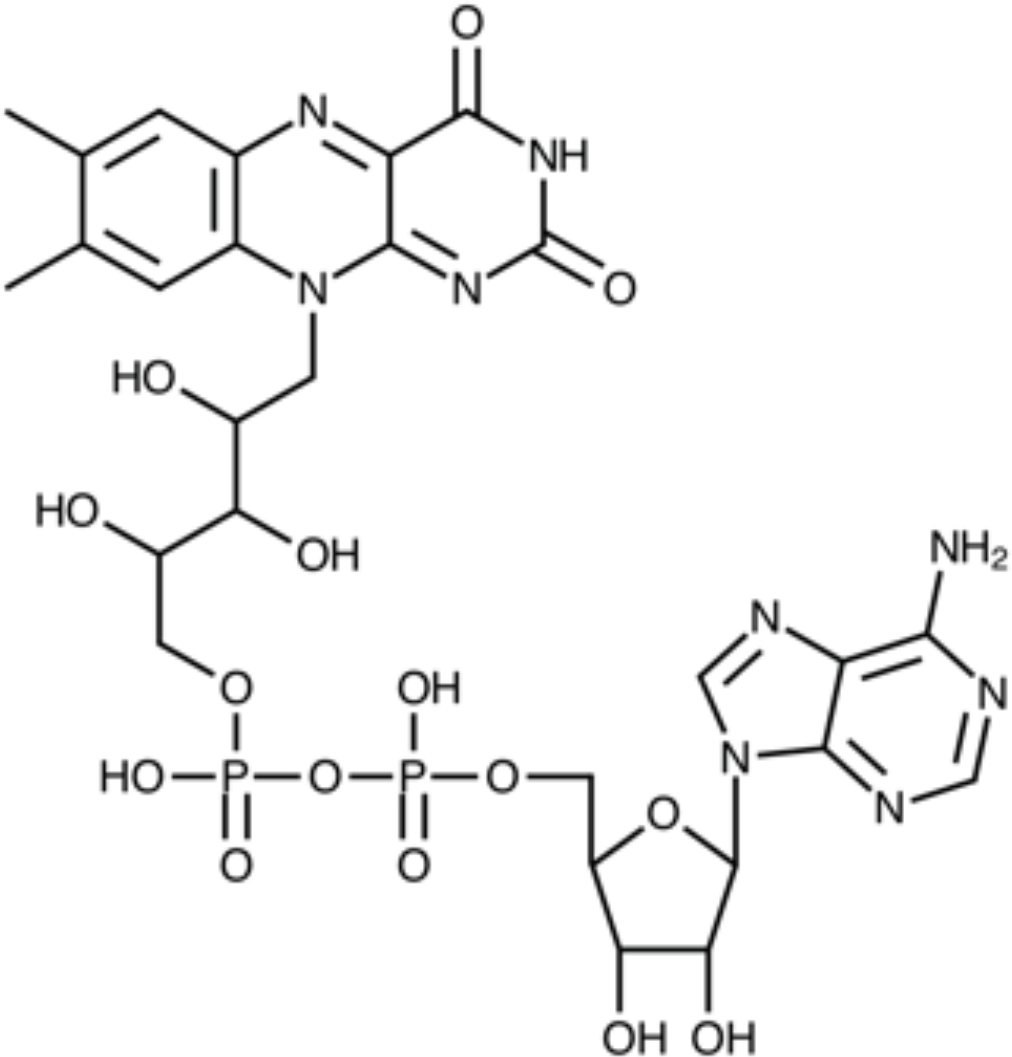
The structural formula of flavin adenine dinucleotide (FAD)

**Figure 3.4.**
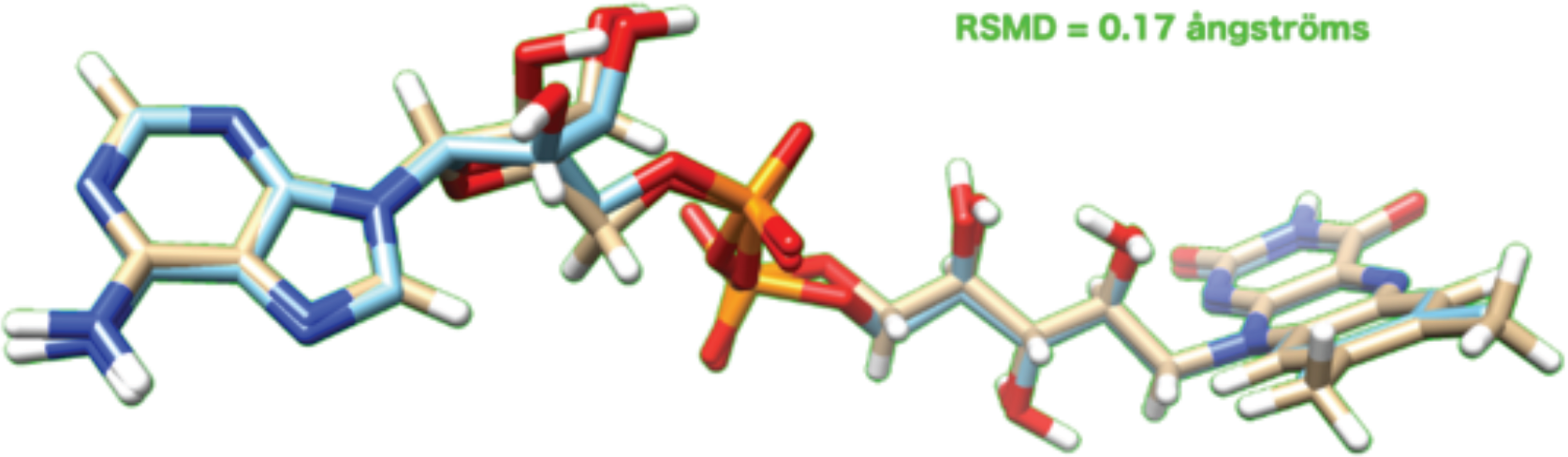
Root mean square deviation of atomic positions between original 2RGH crystal structure FAD and docked FAD in the newly modeled GPD2 cavity.

This position, from now on labeled as the “fully slid in”, was deemed the most stable by the docking calculations.

Three other reasonably stable FAD poses were found by the software, and are summarised, together with their labels and the fully slid in pose, in Table 3.3.

**Table 3.3.** Summary of the reasonably stable FAD poses inside the cavity, as computed by the molecular docking.

| FAD position | FAD binding energy/kcal mol<sup>-1</sup> | Distance from the crystallographic position/Å |
| --- | --- | --- |
| “Fully slid in” | -14.7 | 0 |
| “Slightly slid out” | -12.8 | 3 |
| “Significantly slid out” | -11.9 | 8 |
| “Extremely slid out” | -10.7 | 10 |

The distance from the crystallographic position is reported to quantify the extent of the FAD’s movement.

In Figures 3.5, 3.6, 3.7 and 3.8 the tridimensional representation of the four poses is shown.

**Figure 3.5.**
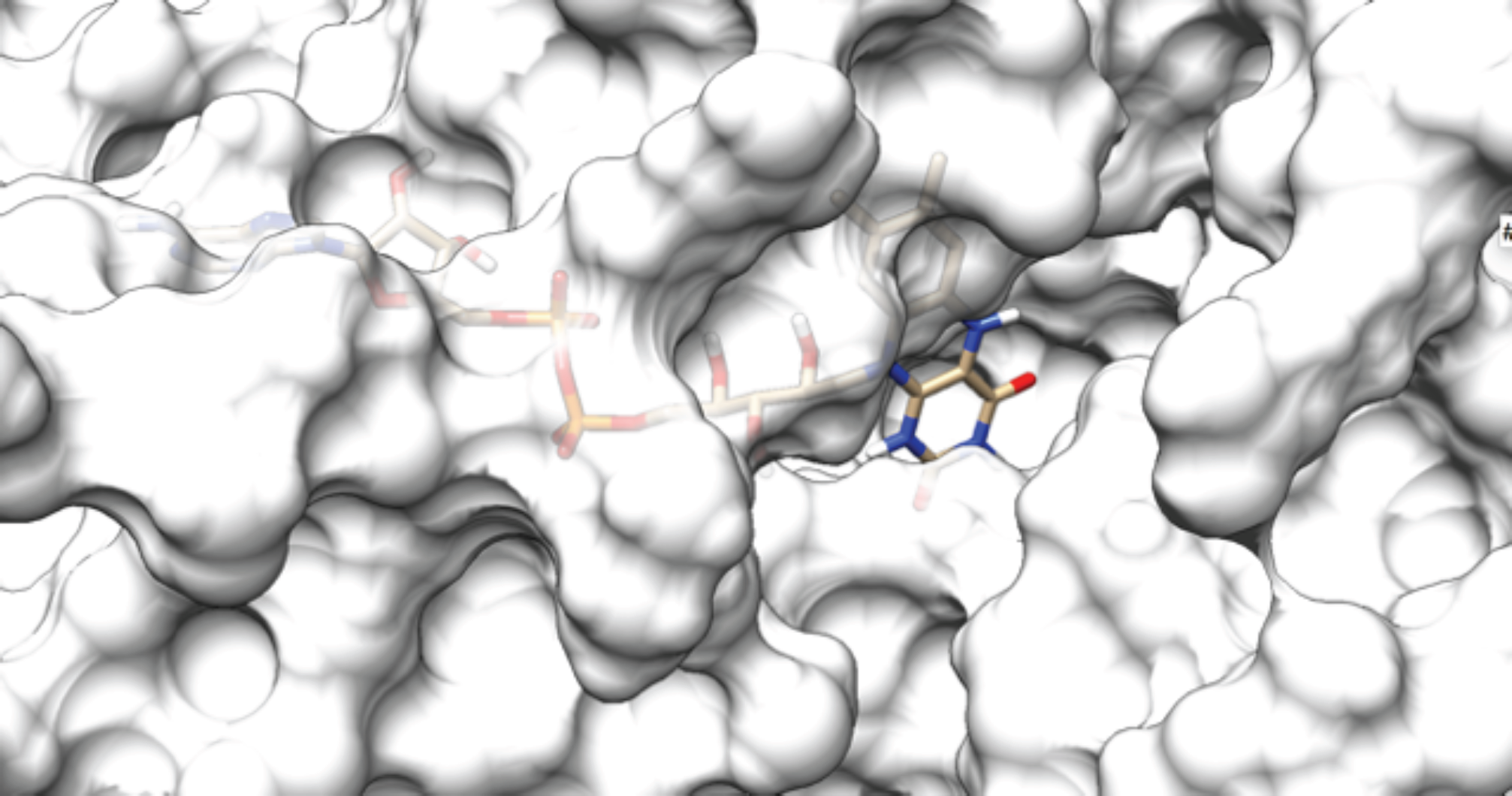
The fully slid in FAD’s pose in the cavity, corresponding to a free binding energy of −14.9 kcal/mol.

**Figure 3.6.**
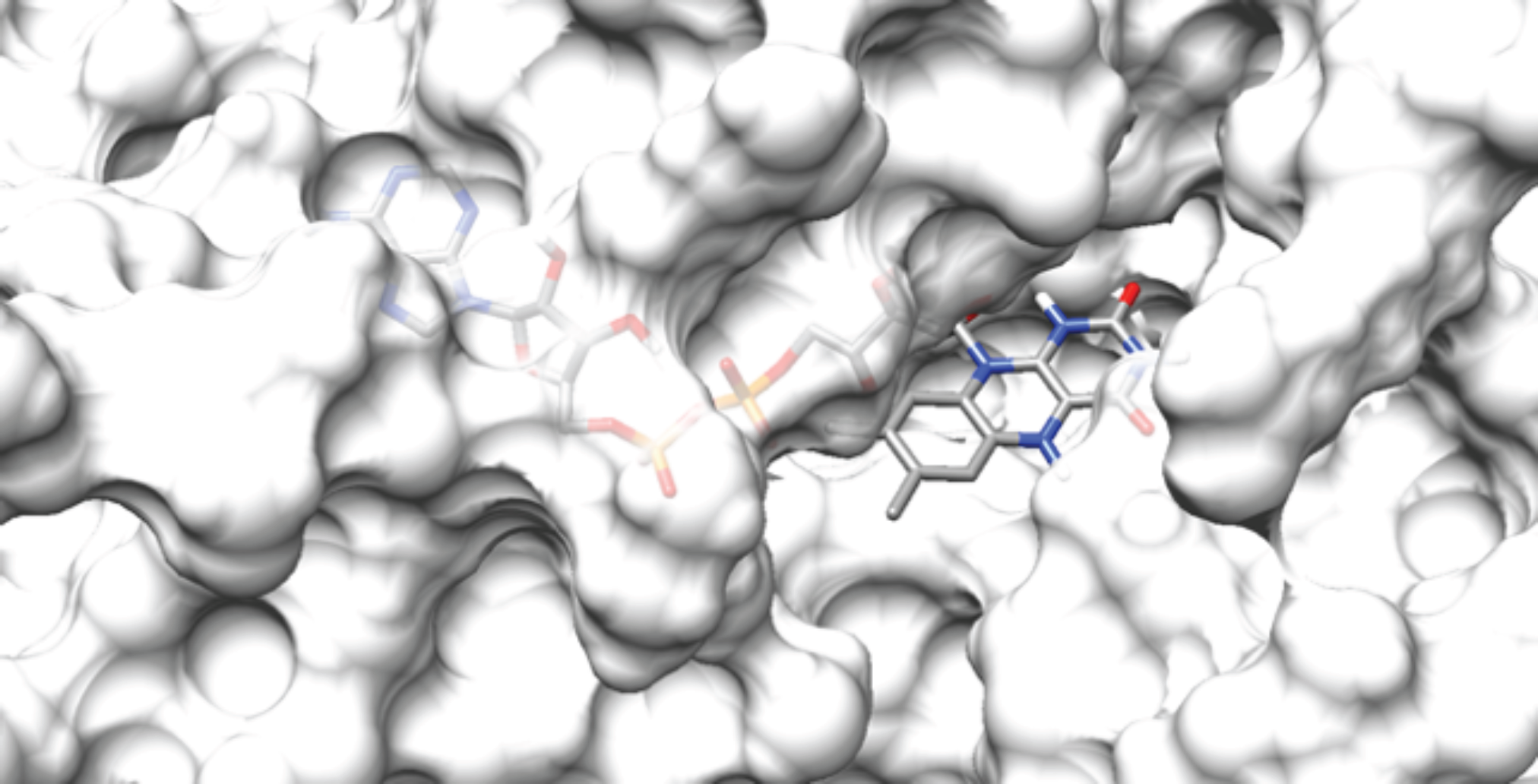
The slightly slid out FAD’s pose in the cavity, corresponding to a free binding energy of −12.6 kcal/mol.

**Figure 3.7.**
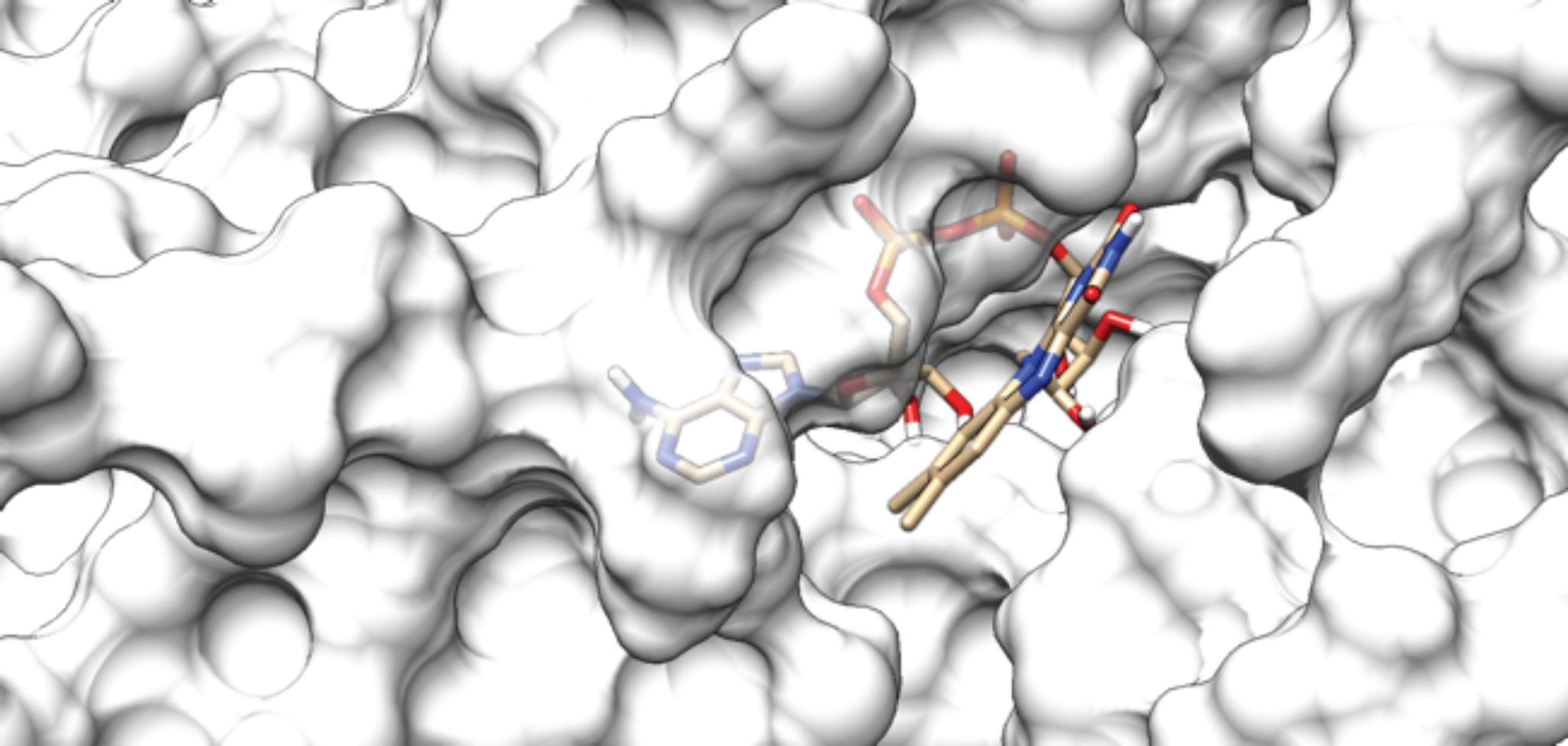
The significantly slid out FAD’s pose, corresponding to a free binding energy of −11.9 kcal/mol.

**Figure 3.8.**
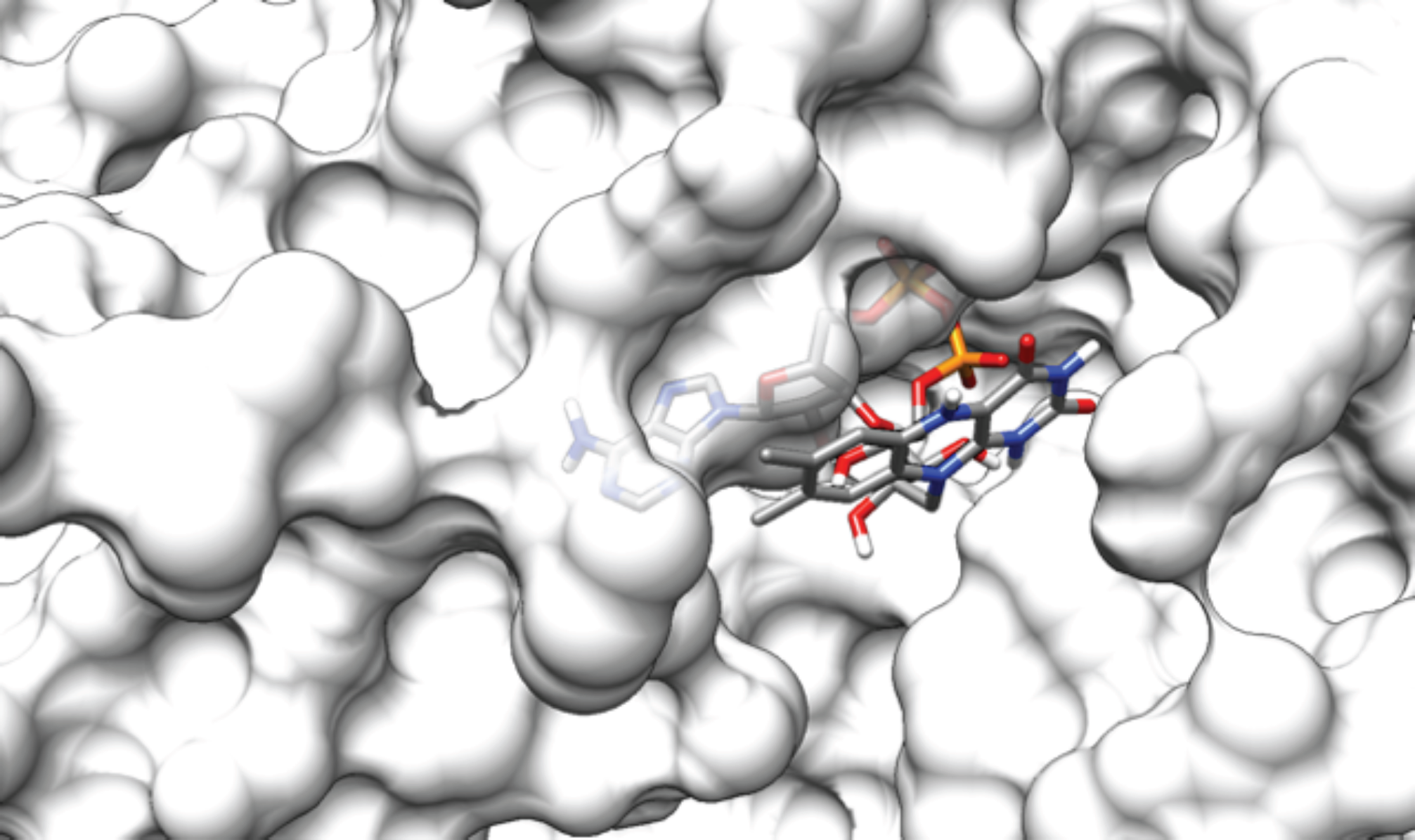
The extremely slid out FAD’s pose in the cavity, corresponding to a free binding energy of −10.6 kcal/mol.

#### 3.2.2 G3P docking

The results of the docking of G3P molecule (Figure 3.9) with the receptor containing the four different FAD poses are summarised in Table 3.4.

**Table 3.4.** Results of the G3P interaction with different FAD poses, as computed by the molecular docking.

| FAD position | G3P binding energy/kcal mol<sup>-1</sup> | Distance between the G3P hydrogen and the FAD nitrogen/Å | Hydrogen bonds | Other bonds (electrostatic, Pi carbon-hydrogen) | Repulsive interactions |
| --- | --- | --- | --- | --- | --- |
| Fully slid in | -6.0 | 2.4 | 4 | 5 | 0 |
| Slightly slid out | -5.1 | 3.4 | 4 | 2 | 1 |
| Significantly slid out | -4.5 | 3.4 | 4 | 1 | 2 |
| Extremely slid out | NA | G3P doesn't bind to FAD | - | - | - |

**Figure 3.9.**
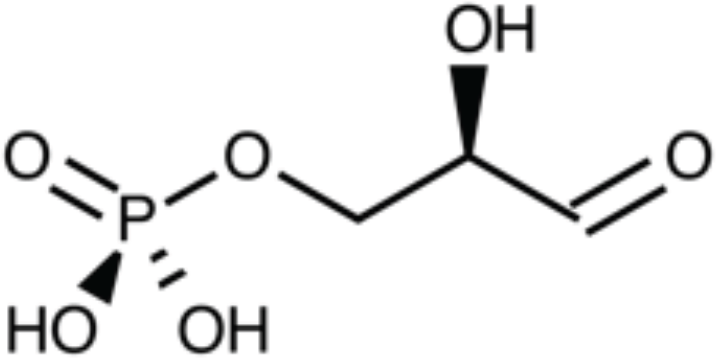
Structural formula of glyceraldehyde 3-phosphate (G3P)

The distance between the G3P’s hydrogen and the FAD’s nitrogen in position 5 on the isoalloxine ring (Figure 3.10) is reported because G3P’s oxidation reaction is believed to initiate from these two atoms (Yeh, Chinte & Du, 2008).

**Figure 3.10.**
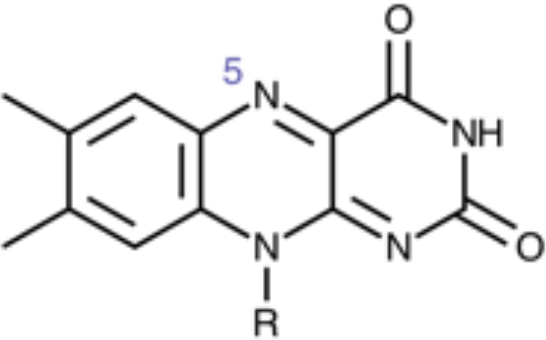
The isoallixine ring of FAD, showing the nitrogen in position 5.

In Figures 3.11-3.13 the cavity interaction of the three G3P docking poses is shown, with highlights on the bond types.

**Figure 3.11.**
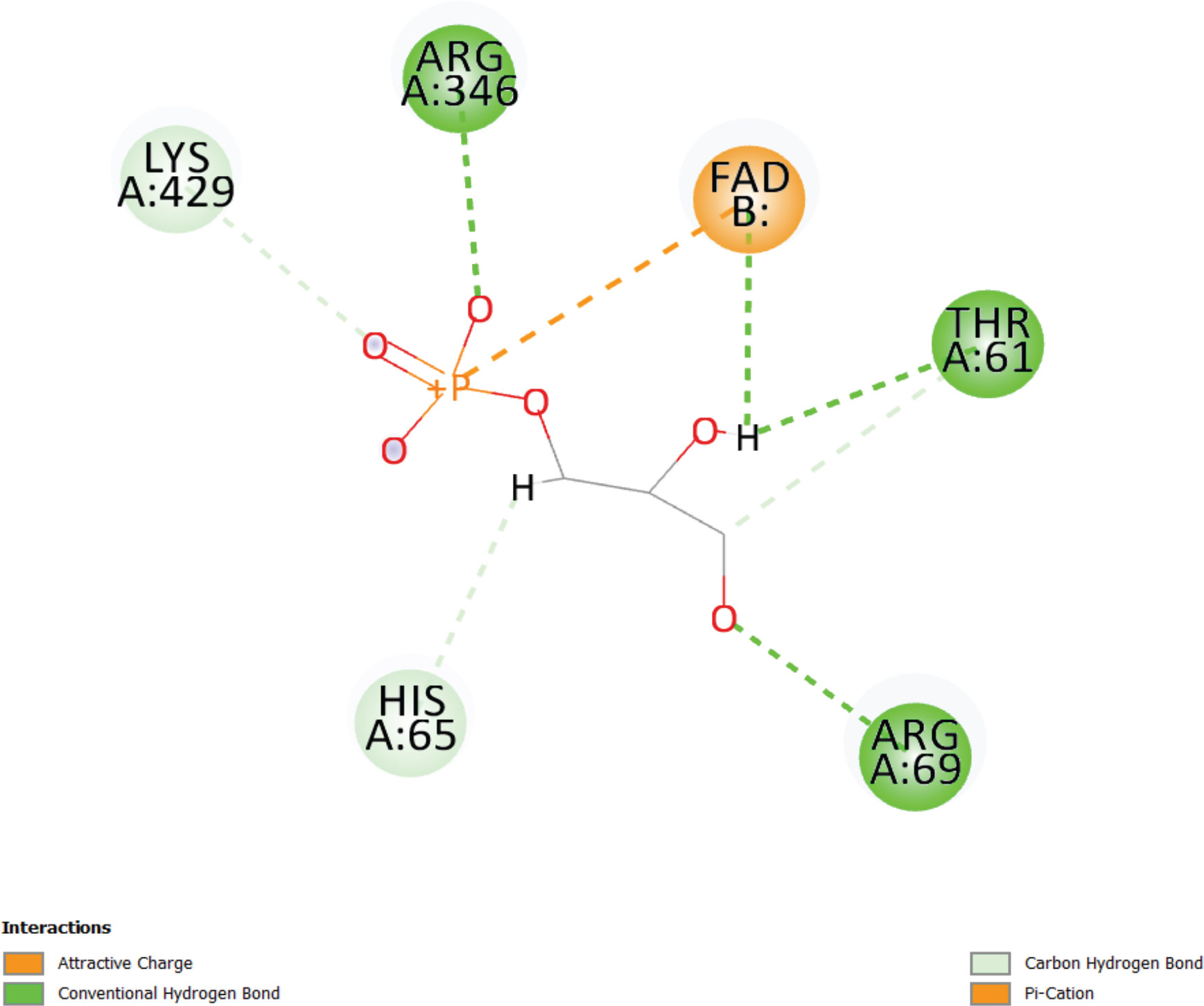
Interaction diagram describes the bonds formed by G3P in the cavity with the fully slid in FAD.

**Figure 3.12.**
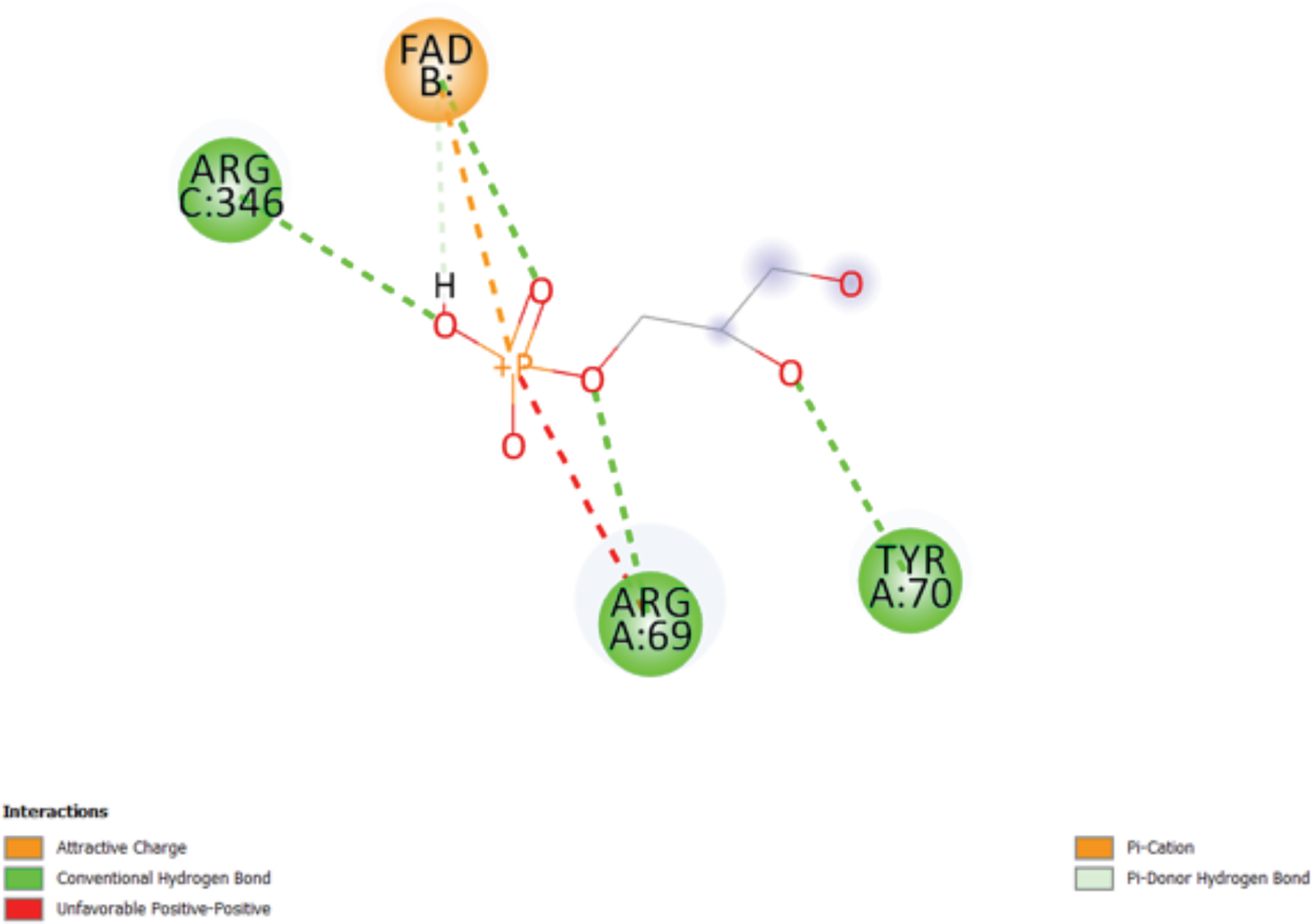
Interaction diagram describes the bonds formed by G3P in the cavity with the slightly slid out FAD.

**Figure 3.13.**
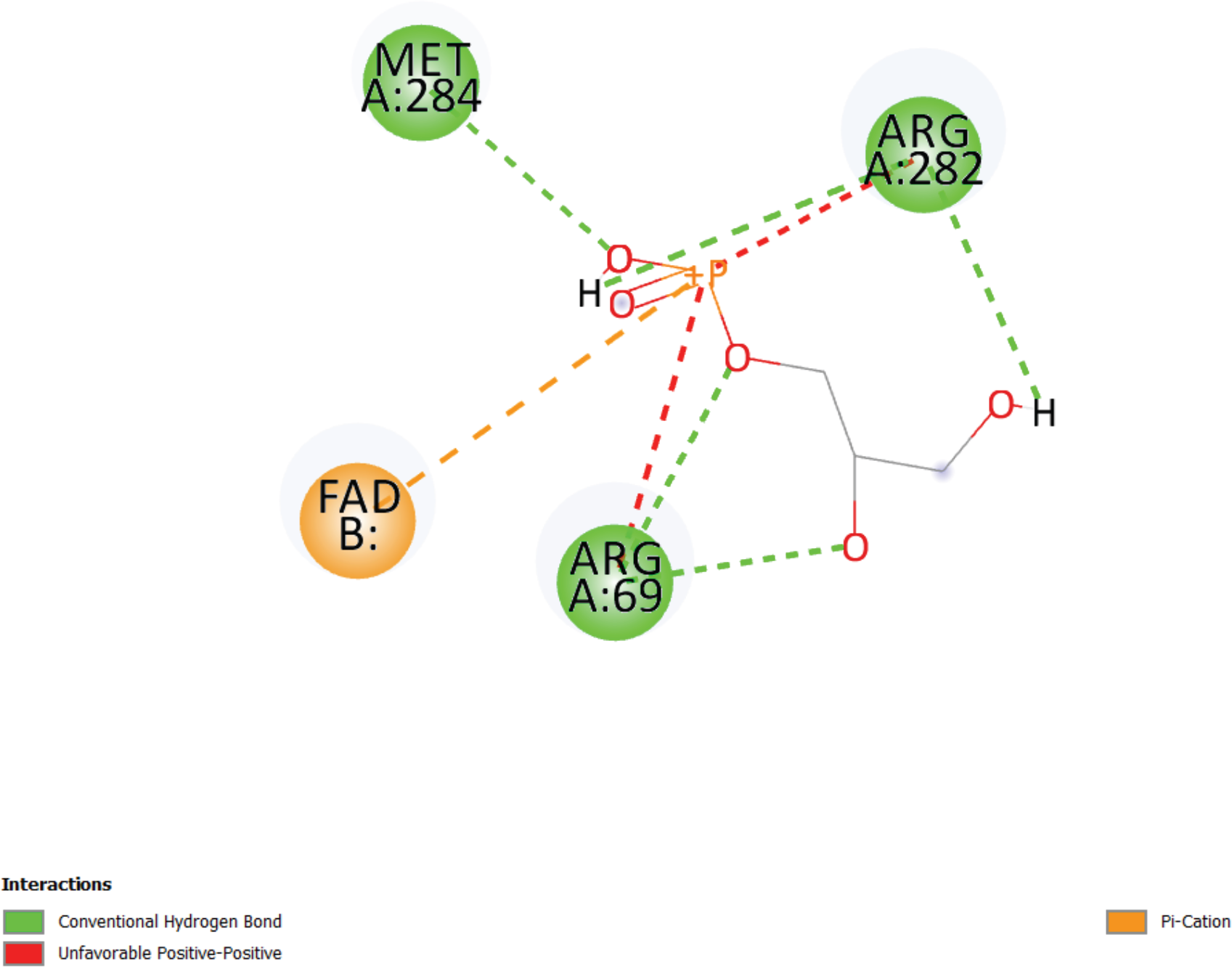
Interaction diagram describes the bonds formed by G3P in the cavity with the significantly slid out FAD.

Overall, G3P showed a decreasing affinity for the binding site with the increasing of the FAD sliding out movement. This agrees with the fact (Fraaije & Mattevi, 2000) that in flavoproteins the docking position of the substrate is buried well below the protein surface to protect the reaction site from the solvent.

#### 3.2.3 FADH2 interactions and redox reactions

In order to assess the most favourable positions for FAD’s reduction and for subsequent FADH_2_ oxidation and electron transfer to ubiquinone, the cavity interactions of the nitrogens in position 1 and 5 on the isoalloxine ring of FAD (Figure 3.14) were analysed for all the four FAD positions.

**Figure 3.14.**
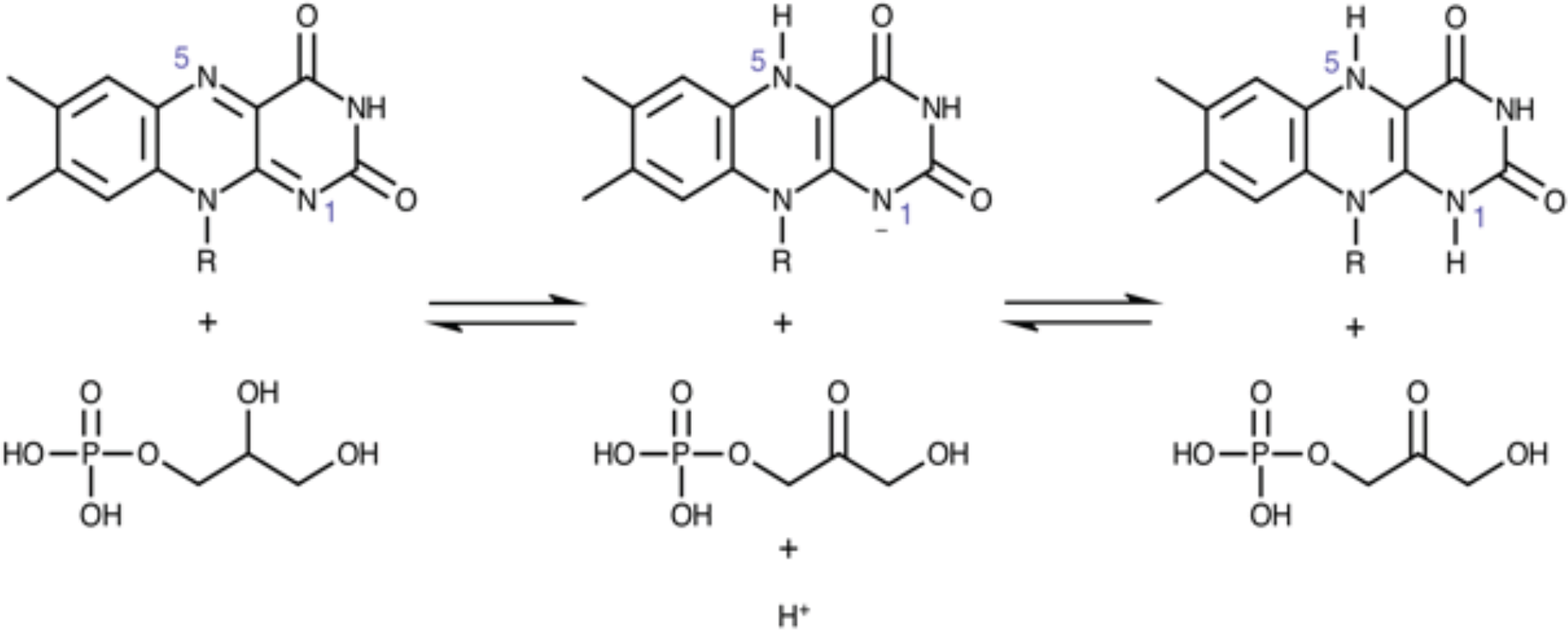
The two steps of the FAD reduction process by glyceraldehyde 3-phosphate (G3P)

According to (Fraaije & Mattevi, 2000) and to (Ghanem & Gadda, 2008), in fact, the FAD in flavoproteins needs a hydrogen bond donor close to position 5 to initiate the reduction reaction, and a positive residue close to position 1 to stabilise the negative charge developed in the intermediate step of the reduction process.

The results are summarised in Table 3.5.

**Table 3.5.** Cavity interactions of the nitrogens in position 1 and 5 on the isoalloxine ring of FADH_2_.

| FAD position | Nitrogen at position 1 | Nitrogen at position 5 |
| --- | --- | --- |
| Fully slid in | No interactions | 2.5 Å hydrogen bond with THR61 |
| Slightly slid out | 3.2 Å proximity with HIS259 | No interactions |
| Significantly slid out | No interactions | No interactions |
| Extremely slid out | No interactions | No interactions |

#### 3.2.4 Coenzyme Q10 docking on GPD2

In order to assess the possibility of a direct electron transfer from FADH_2_ to the final receptor, Coenzyme Q10 (Figure 3.15), the ubiquinone was docked with GPD2 large entrance and its immediate surroundings.

**Figure 3.15.**
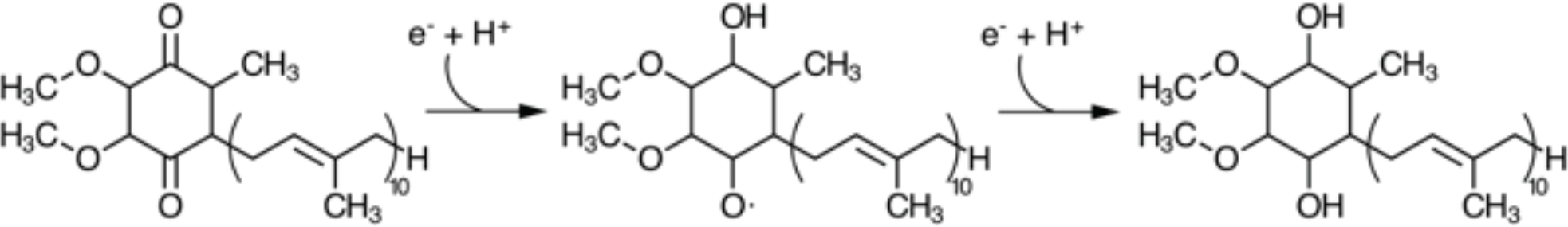
Coenzyme Q10 and its reduction mechanism from left to right.

**Figure 3.16.**
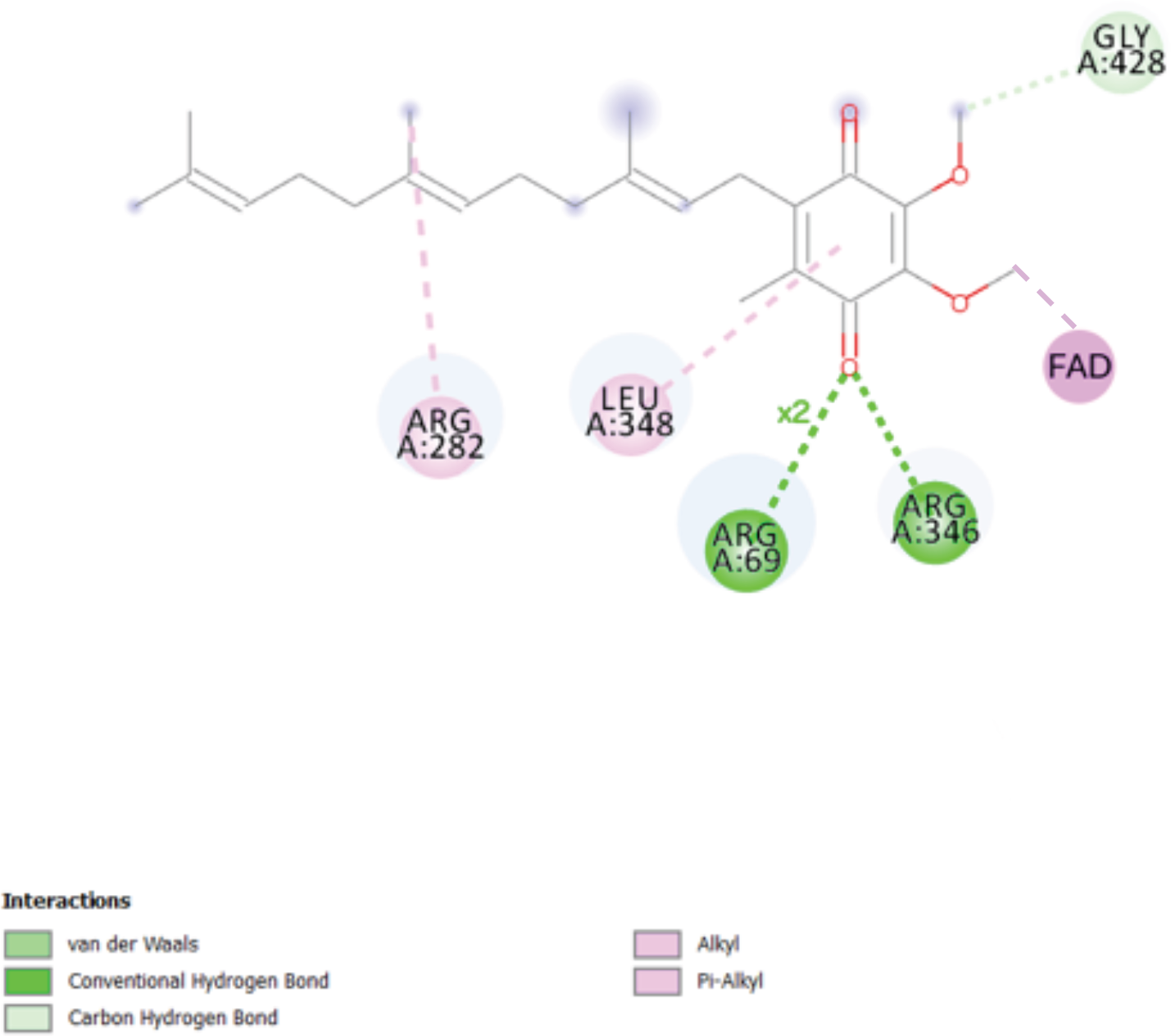
Coenzyme Q10 and its cavity interactions in a 2D representation when FAD is fully slid in.

**Figure 3.17.**
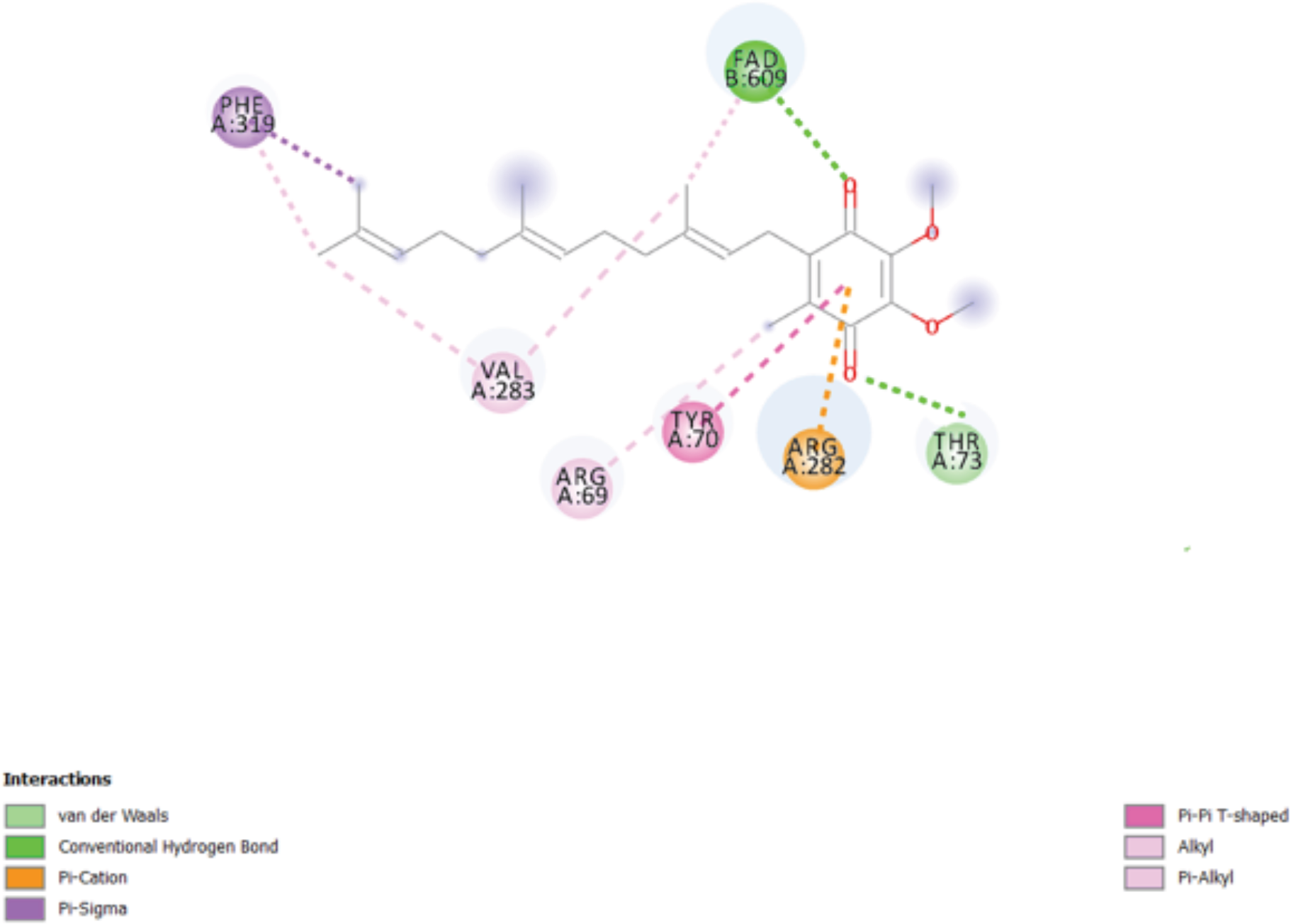
Coenzyme Q10 and its cavity interactions in a 2D representation when FAD is slightly slid out.

**Figure 3.18.**
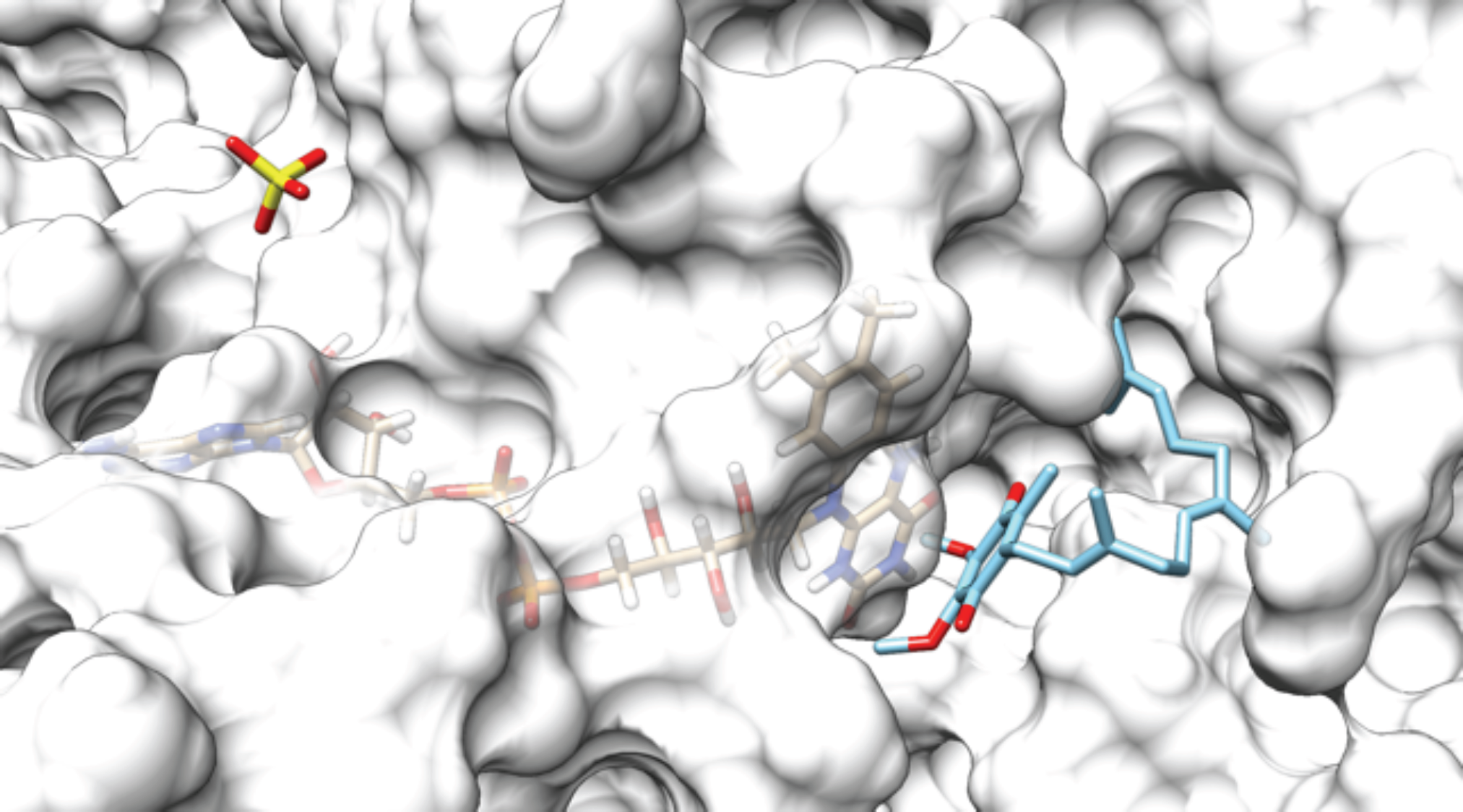
Coenzyme Q10 pose while FAD is fully slid in, having a free binding energy of −6.1 kcal/mol.

**Figure 3.19.**
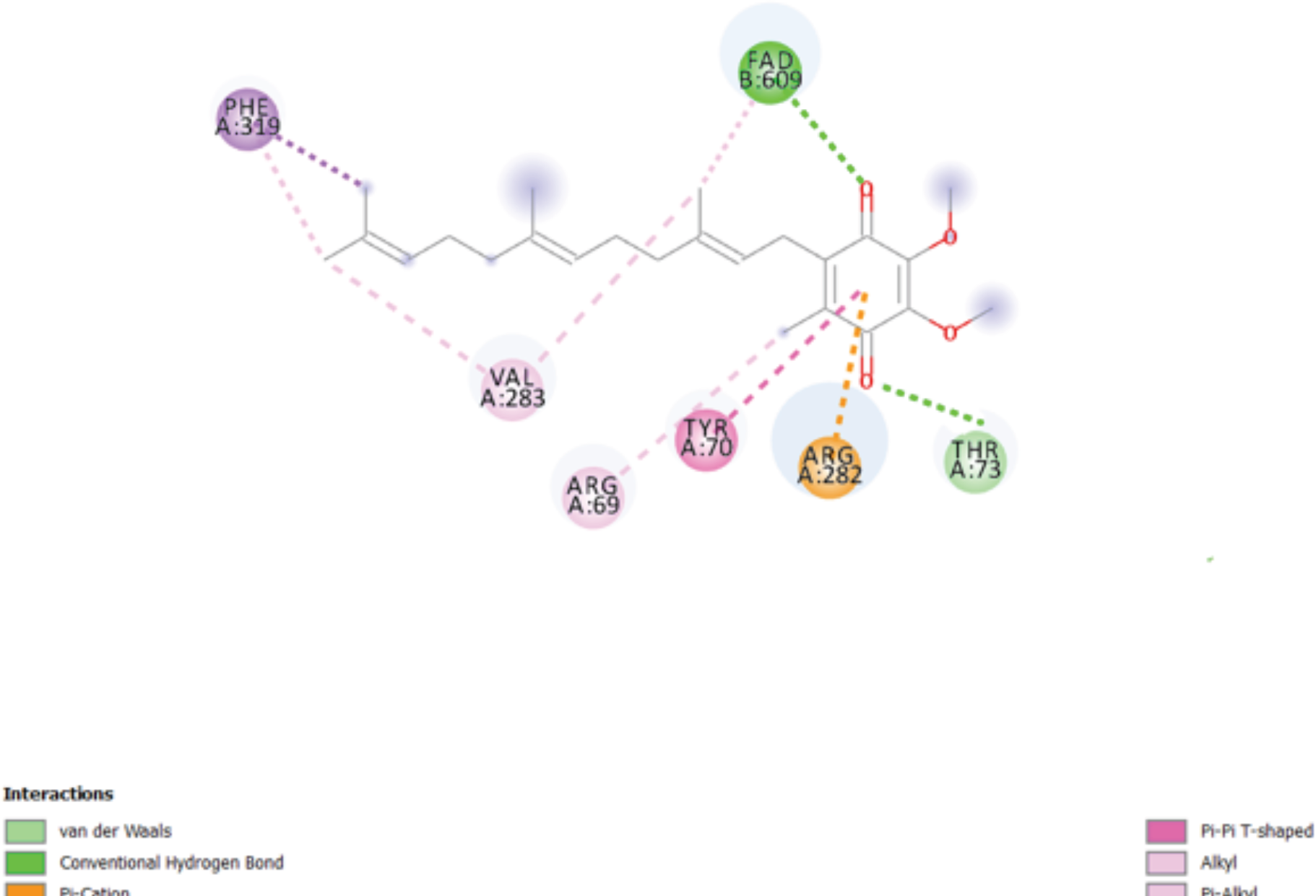
Coenzyme Q10 pose while FAD is slightly slid out, having a free binding energy of −6.6 kcal/mol.

**Figure 3.20.**
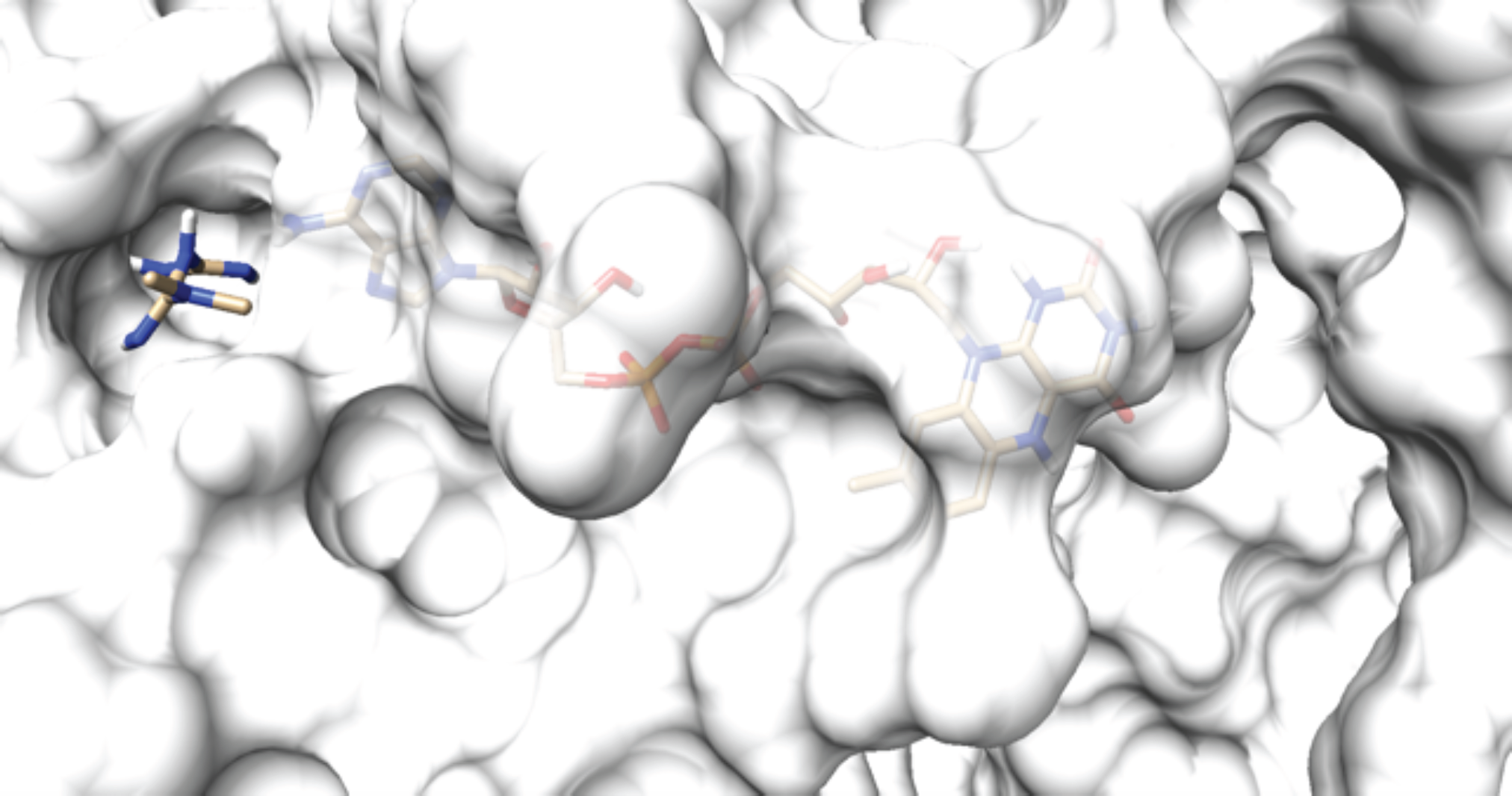
Metformin pose while FAD is slightly slid out, having a free binding energy of −5.0 kcal/mol.

The results of the docking of coenzyme Q10 with the receptor’s large entrance, in correspondance to the four different FAD poses, are summarised in Table 3.6.

**Table 3.6.** Results of the coenzyme Q10 molecular docking with the large entrance with different FAD poses.

| FAD position | Coenzyme Q10 binding energy/kcal mol<sup>-1</sup> | Distance from position 5 nitrogen/ angstroms | Hydrogen bonds | Other bonds |
| --- | --- | --- | --- | --- |
| Fully slid in | -6.1 | 4.7 | 3 | 4 |
| Slightly slid out | -6.6 | 3 | 2 | 8 |
| Significantly slid out | NA - no significative binding | NA - no significative binding | - | - |
| Extremely slid out | NA - no significative binding | NA - no significative binding | - | - |

Coenzyme Q10 shows a good binding ability only when FAD is fully slid in or when it’s slightly slid out. The latter position is slightly preferential from the binding energy point of view, but the former is more specific because it’s characterised by bonds that, on average, are more directional in nature. Finally, the oxidising oxygens of coenzyme Q10 are closer, in the slightly slid out FAD pose, to the target nitrogen in position 5 of the isoalloxine ring. (see Figures 3.21-3.24)

**Figure 3.21.**
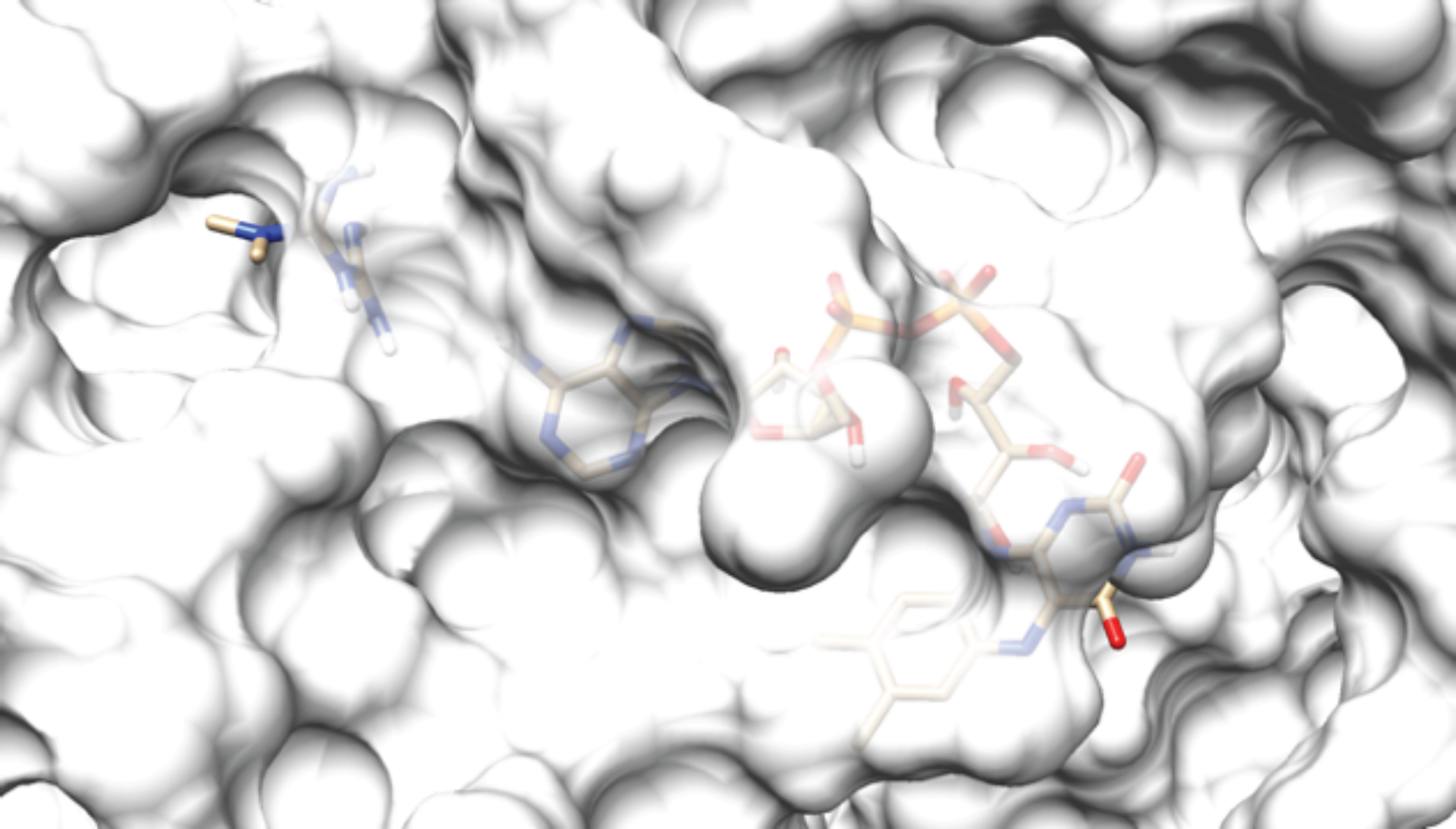
Metformin pose while FAD is significantly slid out, having a free binding energy of −5.3 kcal/mol.

**Figure 3.22.**
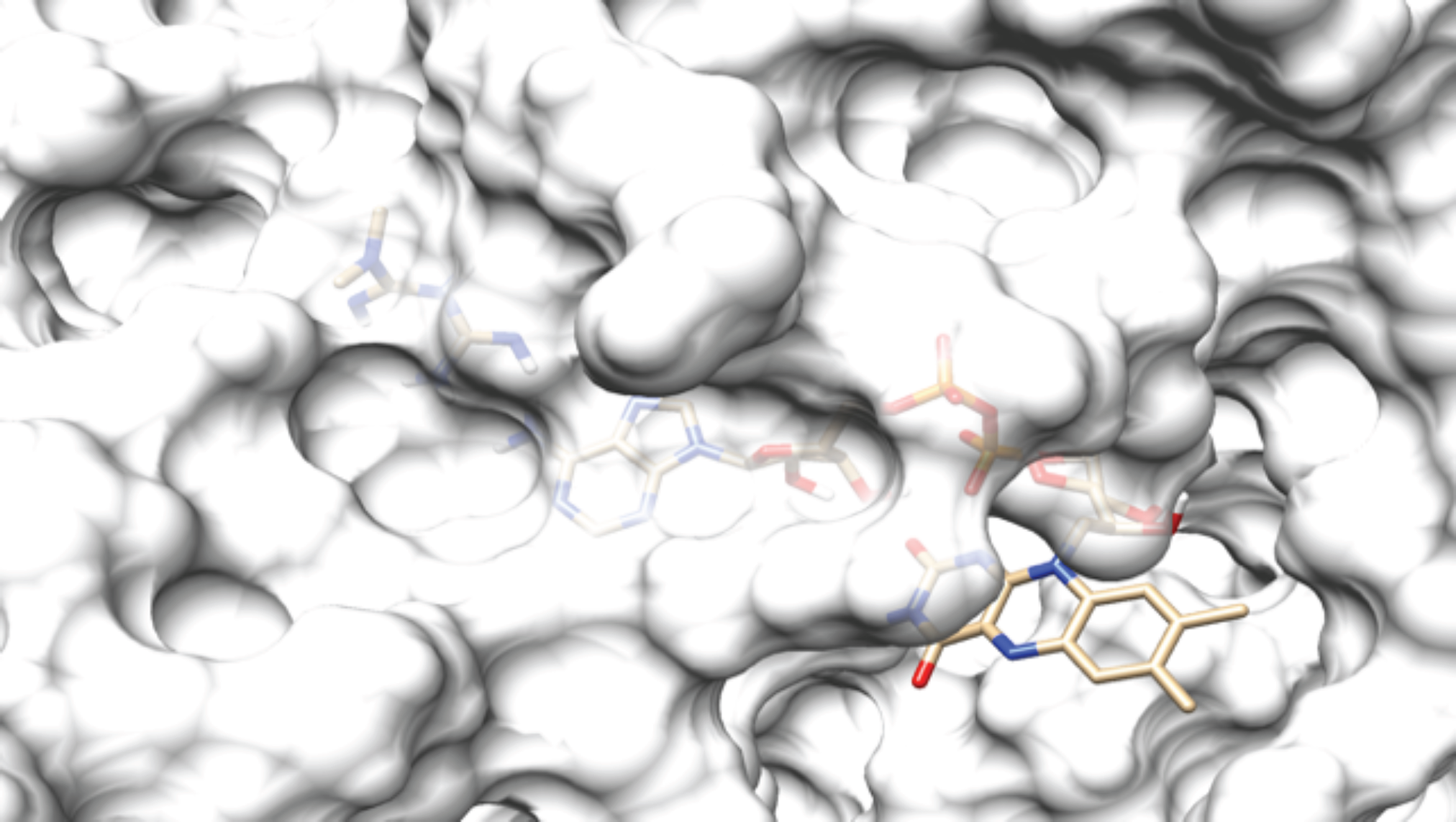
Metformin pose while FAD is extremely slid out, having a free binding energy of −5.8 kcal/mol.

**Figure 3.23.**
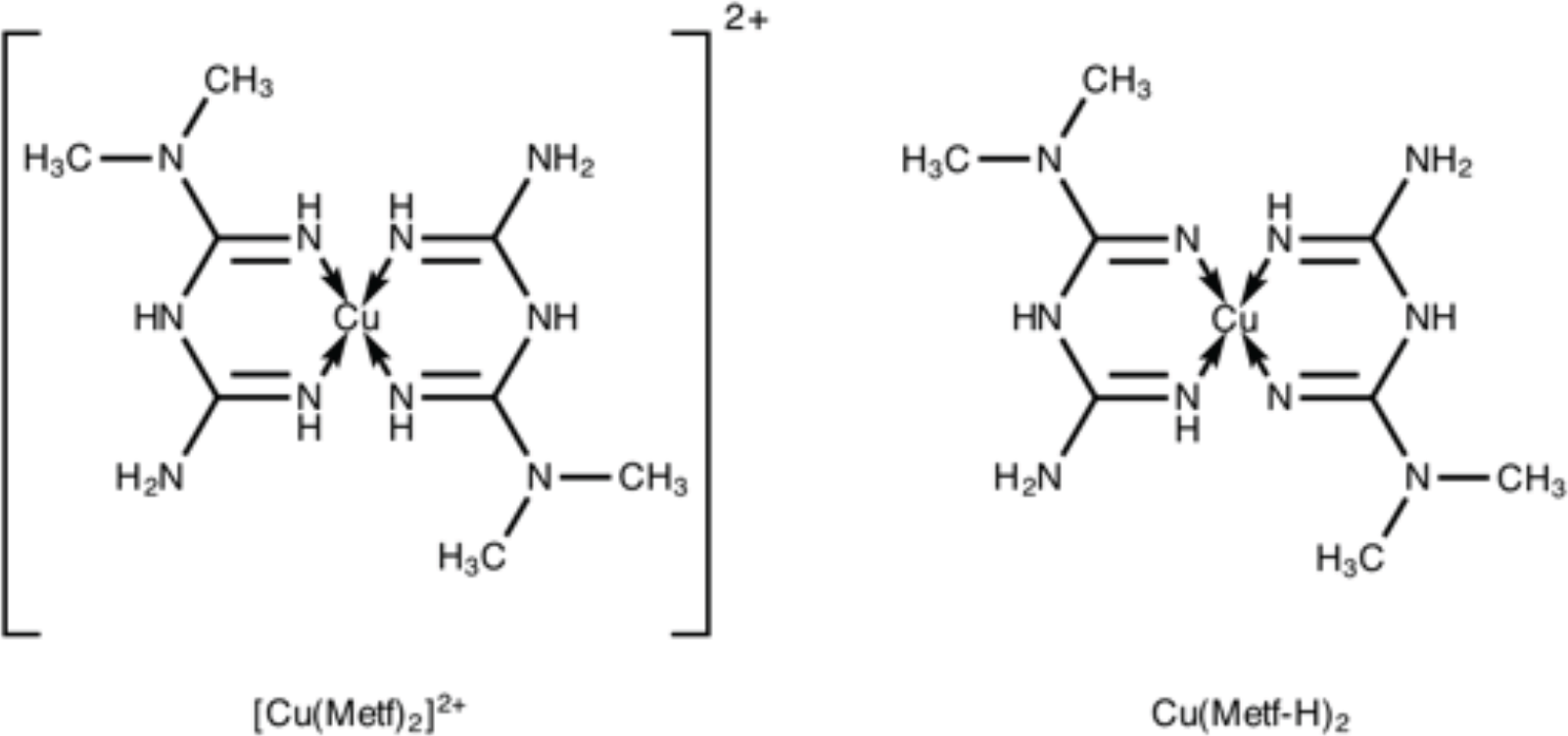
The cationic (left) and neutral (right) dimetformin copper complexes.

**Figure 3.24.**
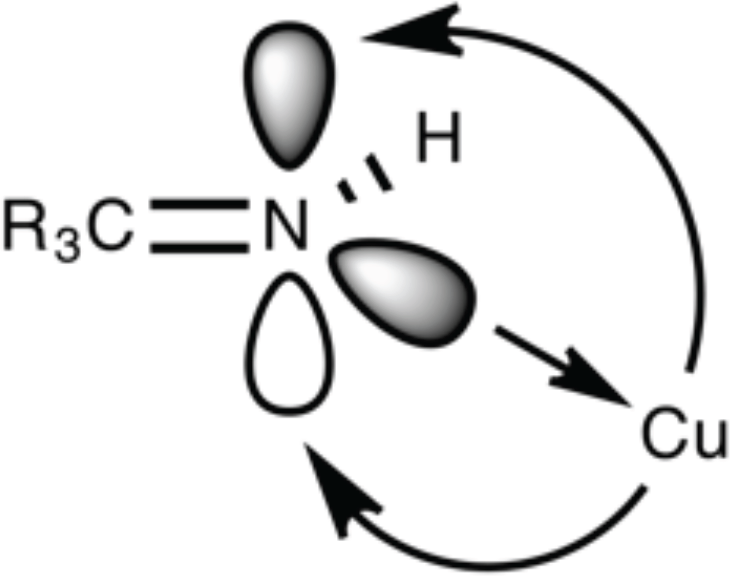
Representation the of the copper coordination through a σ-donation and a π-backdonation.

A mechanism where first the substrate (G3P) docks to the flavine to reduce it and then, after substrate leaves, an ubiquinone docks, in turn, to get reduced, is an example of a so called ping-pong mechanism, something not uncommon in flavoproteins as shown by (Maenpuen et al, 2015).

In Figures 3.16 and 3.17 the 2D interaction diagrams are shown.

In Figures 3.18 and 3.19 the tridimensional representation of the two binding poses is shown.

#### 3.2.5 Metformin at the small entrance

Pure metformin was docked to the small entrance of the FAD cavity in order to assess its ability to block the FAD sliding in movement.

The results of the docking of the metformin molecule with the receptor’s small entrance, in correspondance to the four different FAD poses, are summarised in Table 3.7.

**Table 3.7.** Results of the metformin molecular docking with the small entrance with different FAD poses.

| FAD position | Metformin binding energy/kcal mol<sup>-1</sup> | Metformin inhibition constant(KI)/nM | Hydrogen bonds | Other bonds |
| --- | --- | --- | --- | --- |
| Fully slid in | NA – The small cavity is occupied by FAD | NA – The small cavity is occupied by FAD | - | - |
| Slightly slid out | -5.0 | 2.15 × 10<sup>5</sup> | 2 | 1 |
| Significantly slid out | -5.3 | 1.30 × 10<sup>5</sup> | 1 | 4 |
| Extremely slid out | -5.8 | 5.57 × 10<sup>4</sup> | 3 | 4 |

Metformin doesn’t bind to the small entrance when FAD is fully slid in, and its binding becomes stronger along with FAD’s sliding out movement.

In Figures 3.20, 3.21 and 3.22 the tridimensional representation of the three poses is shown.

#### 3.2.6 Metformin at the large entrance

Pure metformin was docked to the large entrance of the FAD cavity in order to assess its ability to block the FAD sliding out movement.

The results of the docking of metformin molecule with the receptor’s large entrance, in correspondance to the four different FAD poses, are summarised in Table 3.8.

**Table 3.8.**
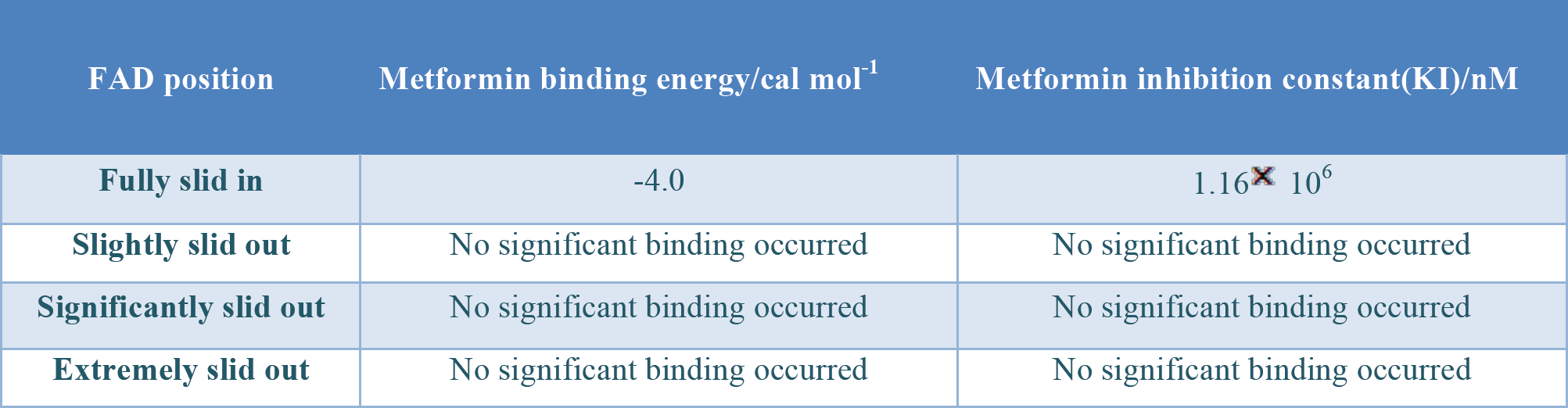
Results of the metformin molecular docking with the large entrance with different FAD poses.

| FAD position | Metformin binding energy/cal mol<sup>-1</sup> | Metformin inhibition constant(KI)/nM |
| --- | --- | --- |
| Fully slid in | -4.0 | 1.16 × 10<sup>6</sup> |
| Slightly slid out | No significant binding occurred | No significant binding occurred |
| Significantly slid out | No significant binding occurred | No significant binding occurred |
| Extremely slid out | No significant binding occurred | No significant binding occurred |

All metformin binding poses at the large entrance resulted either far from FAD or with free binding energies not better than −4.0 kcal/mol.

#### 3.2.7 Dimetformin copper complex at the large entrance

In order to assess their ability to block the FAD sliding movement or to block the coenzyme Q10 docking, both the cationic and the neutral metformin copper complexes (Figure 3.23) were docked to the small and to the large entrance of the FAD cavity.

The planar pseudo-aromatic structures of the complexes were realised using the (Repiščák et al., 2014) hypothesis of a metal coordination through a σ-donation and a π-backdonation with a delocalised π ring system being formed (Figure 3.24).

They both showed affinity with the large entrance of the binding cavity, while they were unable to dock to the small entrance.

The results of the docking of the copper complexes with the receptor’s large entrance, in correspondance to the full slid in FAD pose, are summarised in Table 3.9.

**Table 3.9.**
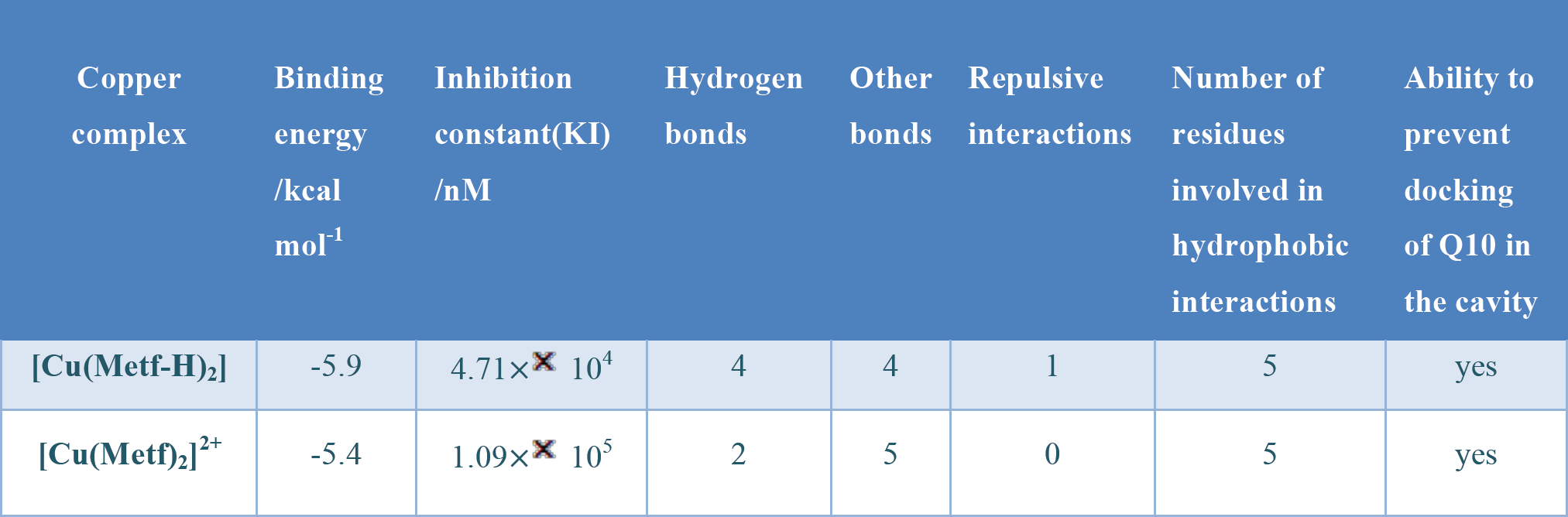
Results of the dimetformin copper complex molecular docking with the large entrance with different FAD poses.

The detailed interaction of the neutral complex with the cavity residues is summarised in Figure 3.25.

**Figure 3.25.**
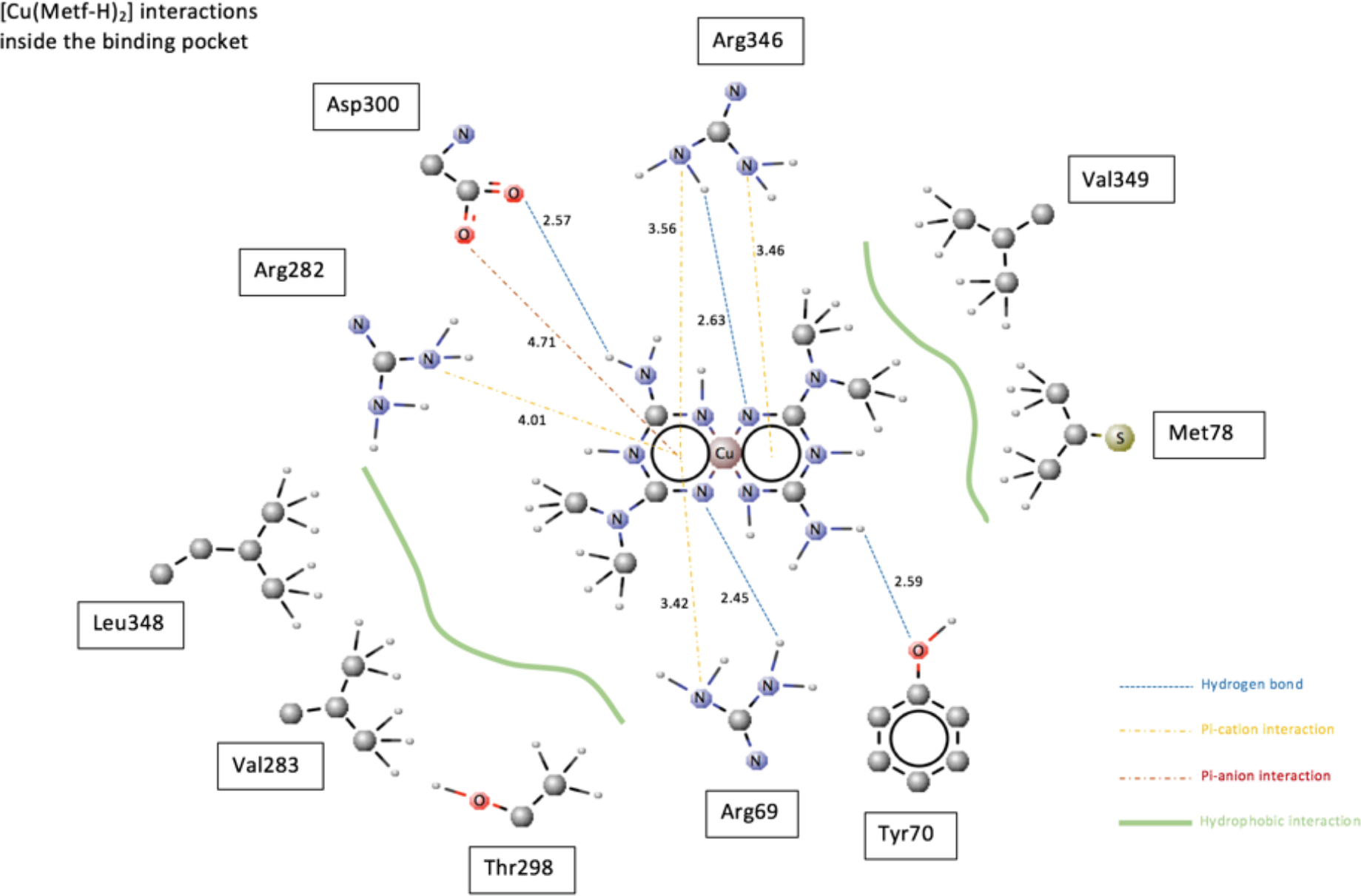
[Cu(Metf-H)_2_] interactions with cavity residues, bond lengths are expressed in angstroms.

It’s worth noticing two facts about the above figure:

1. the interactions of the cationic complex are very similar yet less effective, with the Pi-anion repulsion substituted by a charge-charge attraction with the oxygen of ASP300, and two missing hydrogen bonds caused by the fact that the two hydrogen bond acceptors of the neutral complex are substituted by the protonated nitrogens of the cationic form
2. the Pi-anion repulsion characterising the neutral complex pose has an indirect stabilising effect on the pose, because it “pushes away” the negative aromatic cloud from the ring allowing it to be involved more strongly in the three Pi-cation interactions

The tridimensional representation of the neutral copper complex position relative to the interacting residues is shown in Figure 3.26.

**Figure 3.26.**
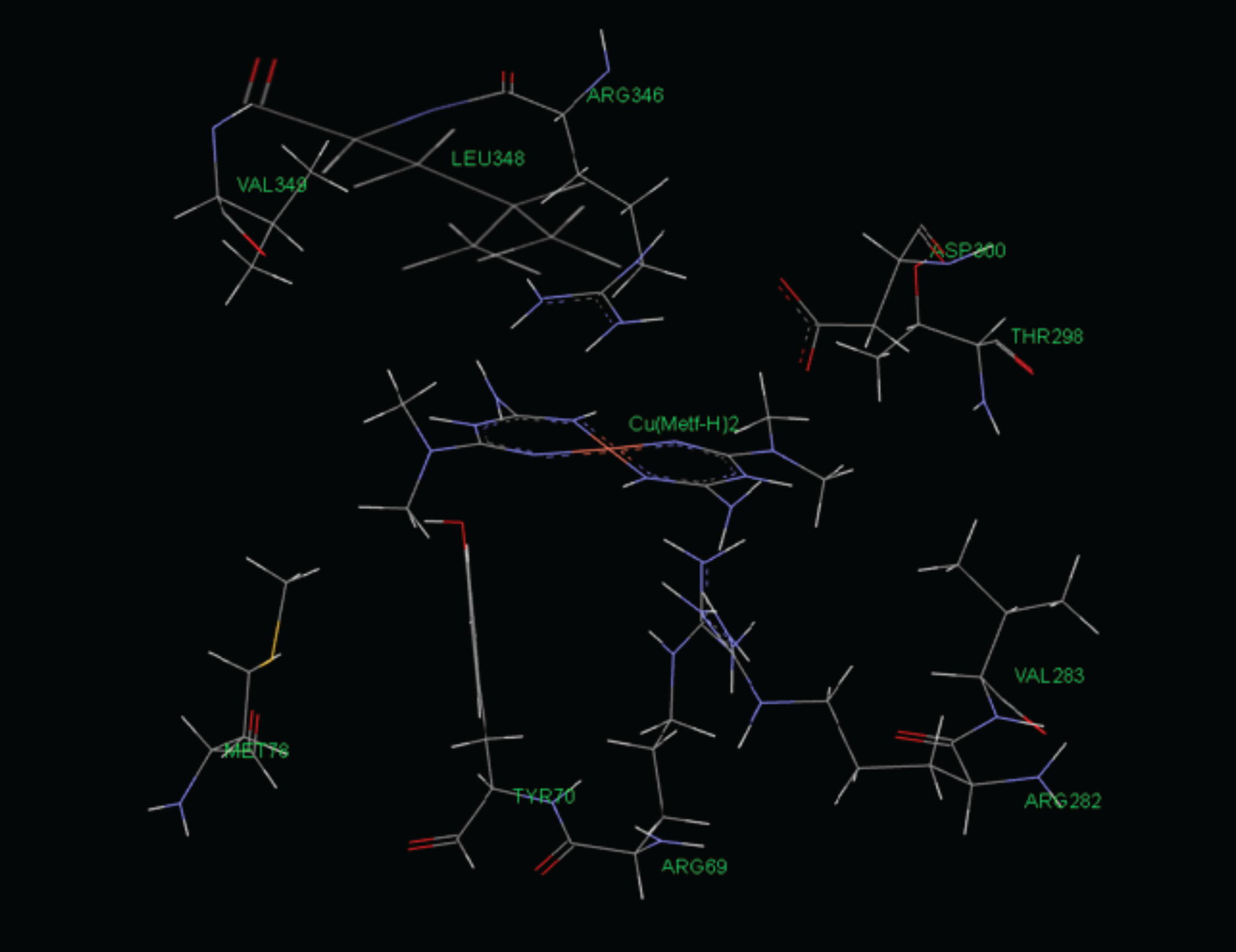
[Cu(Metf-H)_2_] position relative to cavity residues.

When the pose of the dimetformin copper complex docked with the fully slid in FAD and the pose of the slightly slid out FAD are superimposed a collision is shown, demonstrating that the copper complex is able to block the FAD sliding out movement. This is shown in Figure 3.27.

**Figure 3.27.**
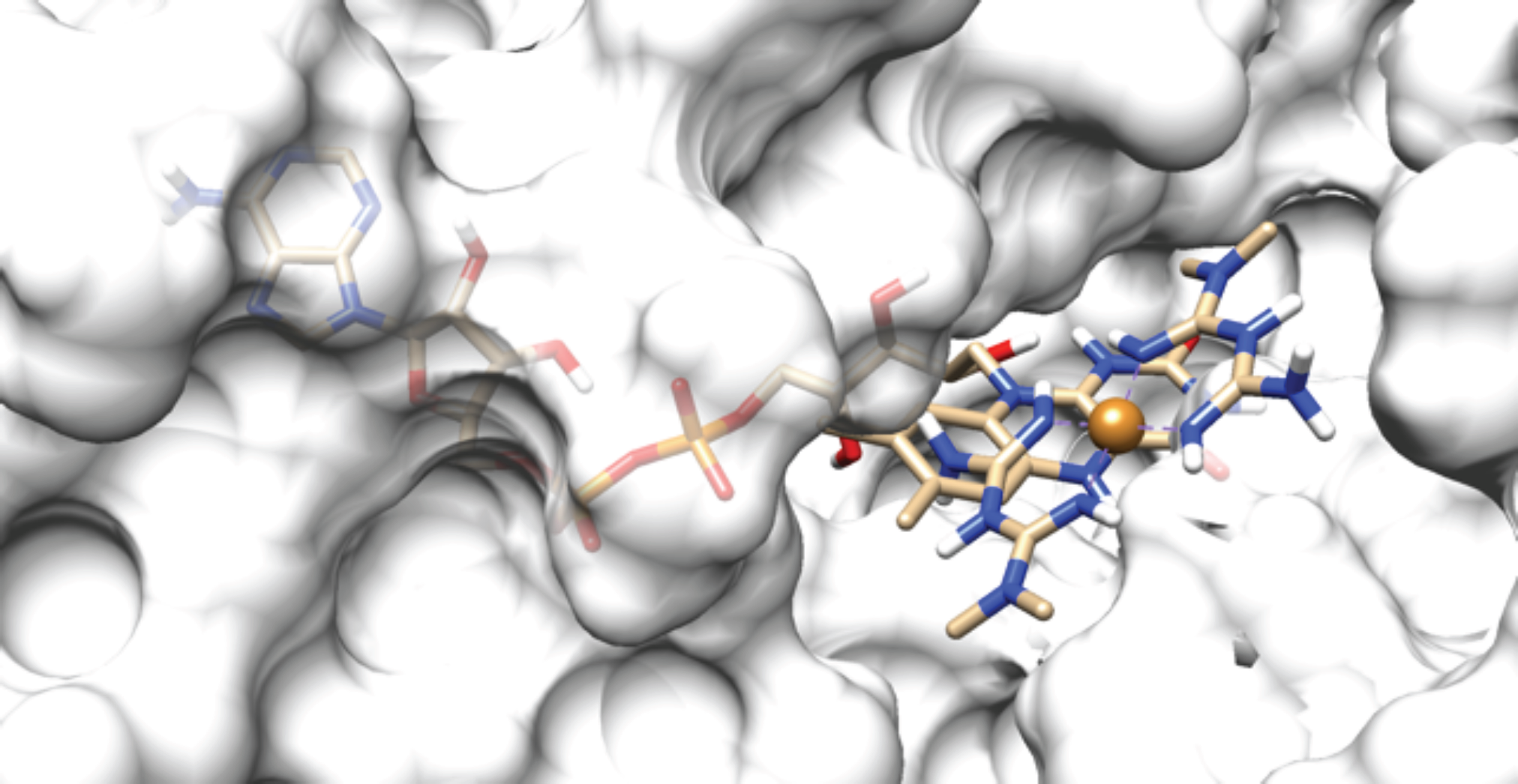
[Cu(Metf-H)_2_] collision with slightly slid out FAD, GPD2 is shown as a 40% transparent HF surface.

The more protruding FAD positions are blocked as well.

Finally, a docking simulation was performed, docking G3P to the cavity’s large entrance when the dimetformin copper complex was bound too, showing that G3P docks to GPD2, and close to the FAD’s head, with the same poses and binding energies no matter if the copper complex is present or not, and therefore the complex satisfies the requisites of a non-competitive inhibitor.

In Figures 3.28 and 3.29 this “cohabitation” between G3P and the complex is shown.

**Figure 3.28.**
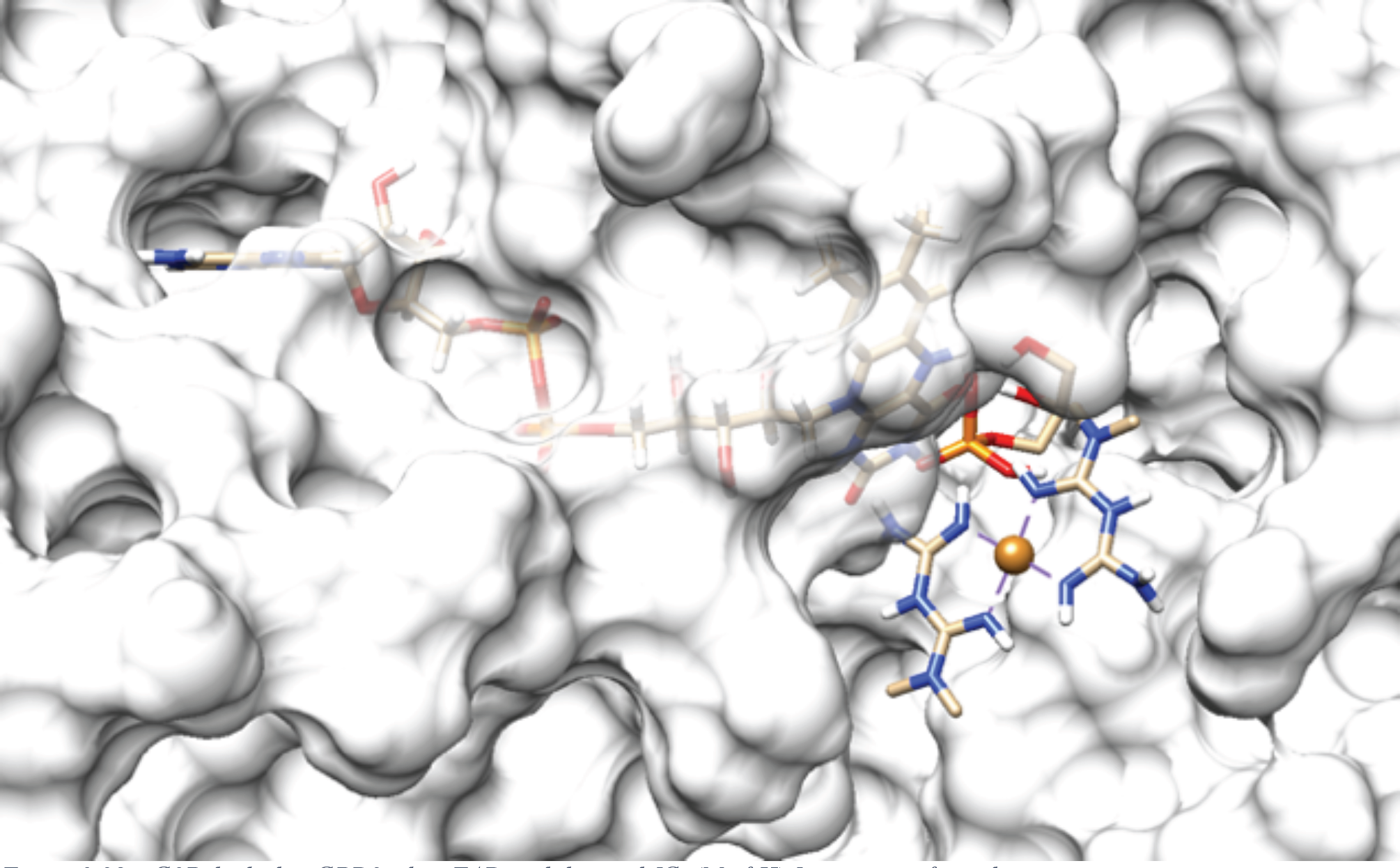
G3P docked to GPD2 when FAD is slid in and [Cu(Metf-H)_2_] is present, frontal view.

**Figure 3.29.**
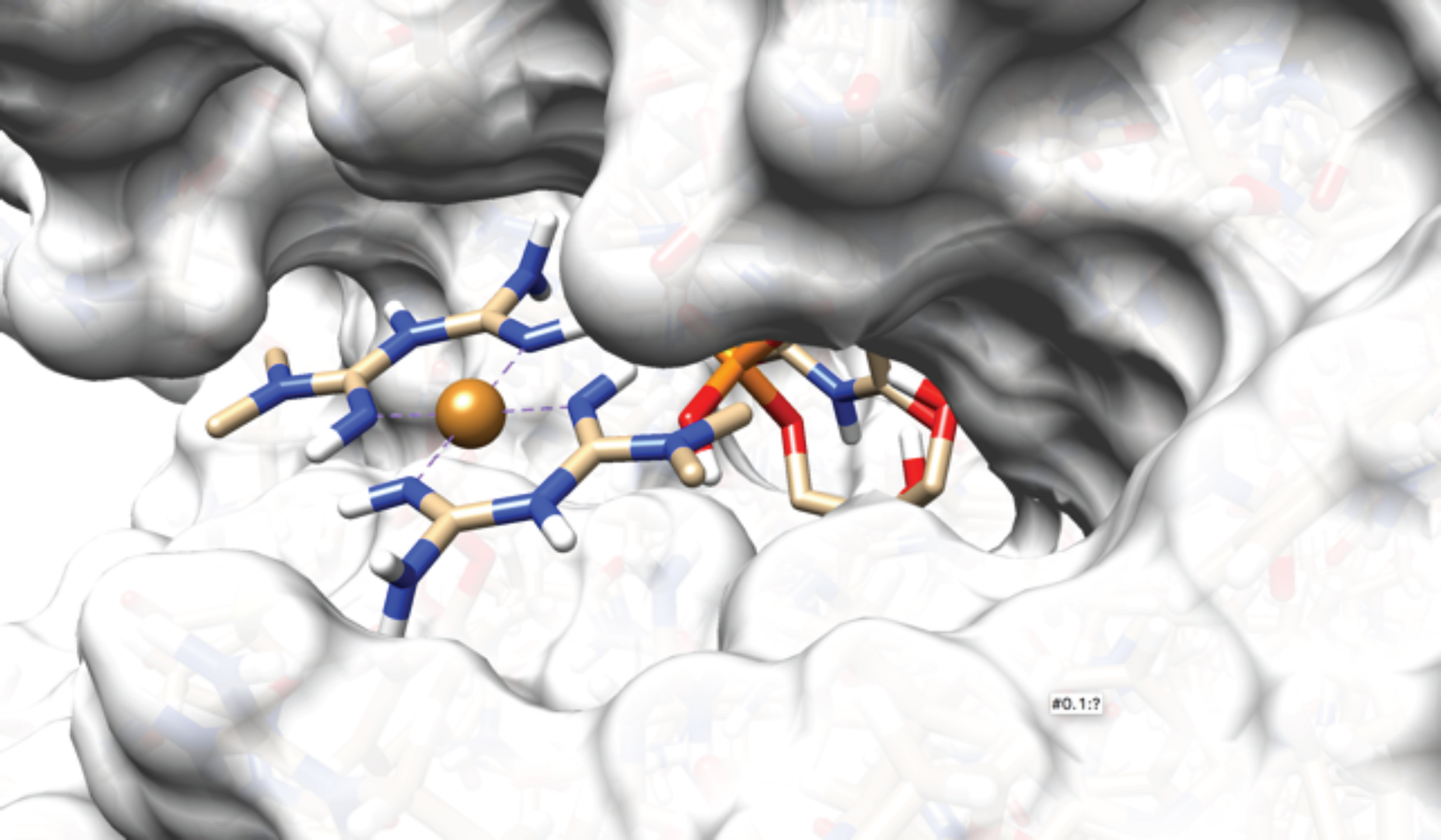
G3P docked to GPD2 when FAD is slid in and [Cu(Metf-H)_2_] is present, lateral view.

#### 3.2.8 Phenformin docking at the small and at the large entrance

Pure phenformin (Figure 3.30) was docked to both the small and the large entrance of the FAD cavity, in analogy of what was done with metformin in paragraphs 3.2.5 and 3.2.6 above.

**Figure 3.30.**
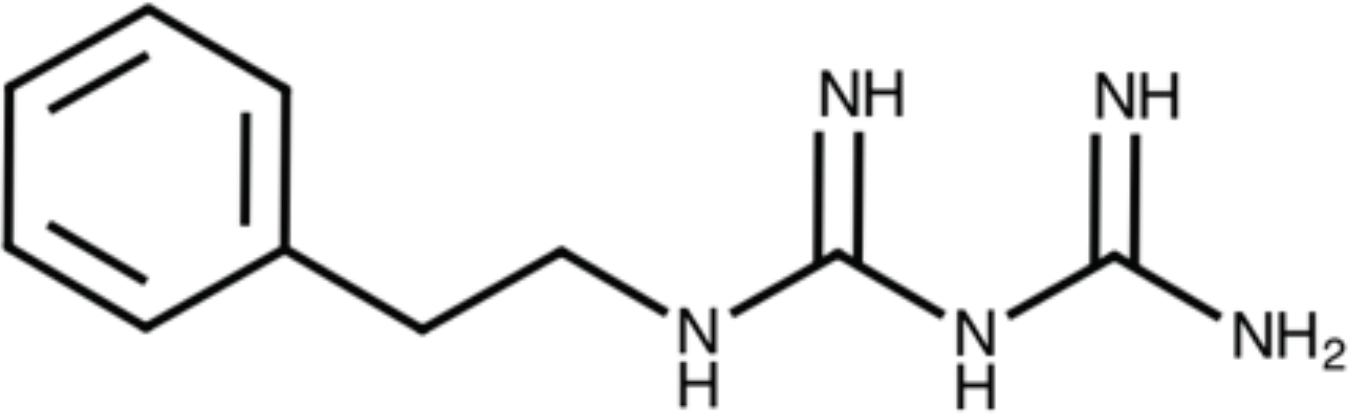
The structural formula of phenformin.

The results are similar to the ones of metformin, with phenformin showing no significant binding to the large exit and with a binding to the small exit becoming stronger along with FAD’s sliding out movement, with better binding energies than metformin, as summarised in Table 3.9.

In Figure 3.31 phenformin binding to the cavity’s small entrance when FAD is slightly slid out is shown.

**Figure 3.31.**
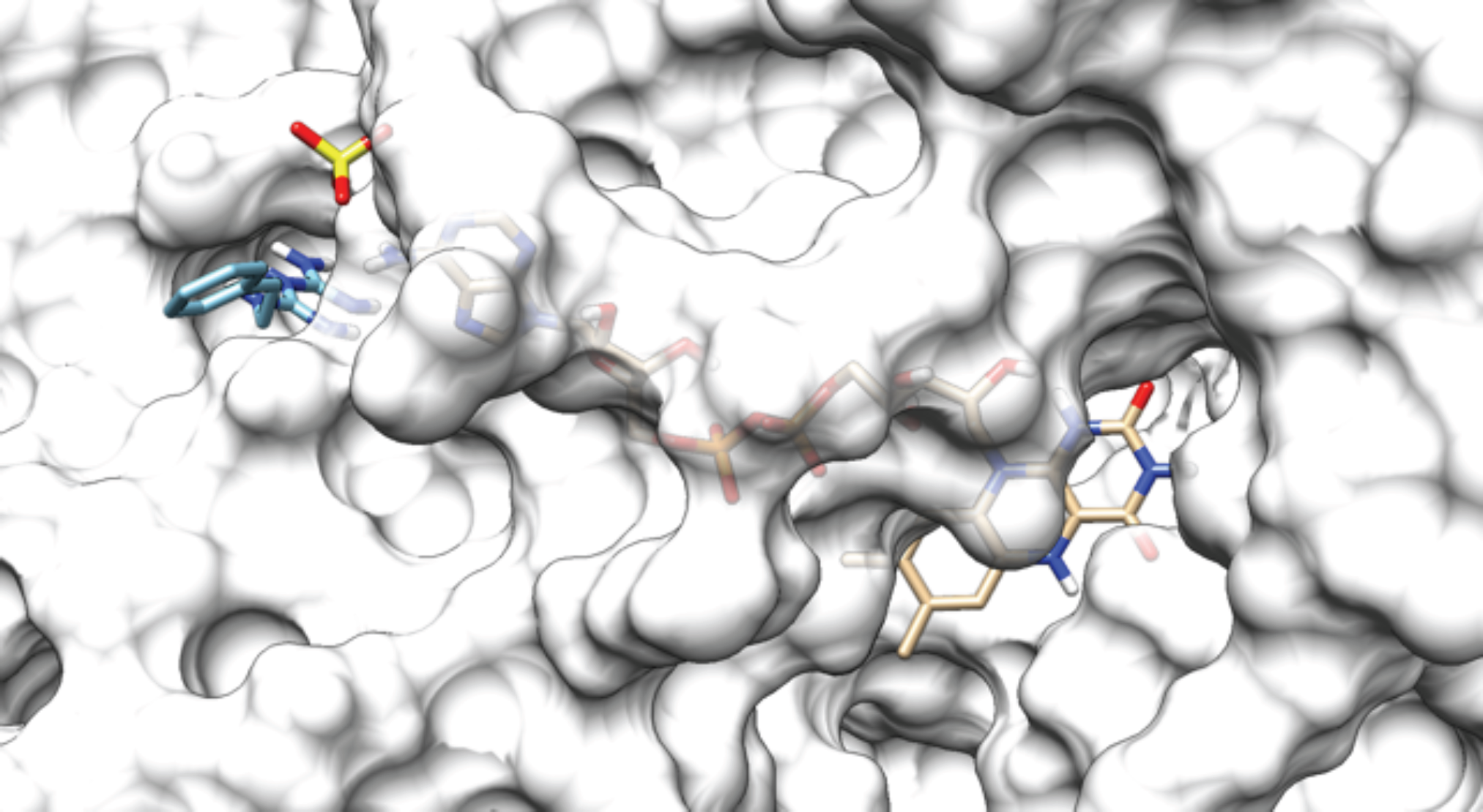
Phenformin binding to the cavity’s small entrance when the FAD is slightly slid out, corresponding to a binding energy of −5.7 kcal/mol.

#### 3.2.9 Monophenformin copper complex at the large entrance

The monophenformin copper complex (Figure 3.32) was docked to both the small and the large entrance of the FAD cavity.

**Figure 3.32.**
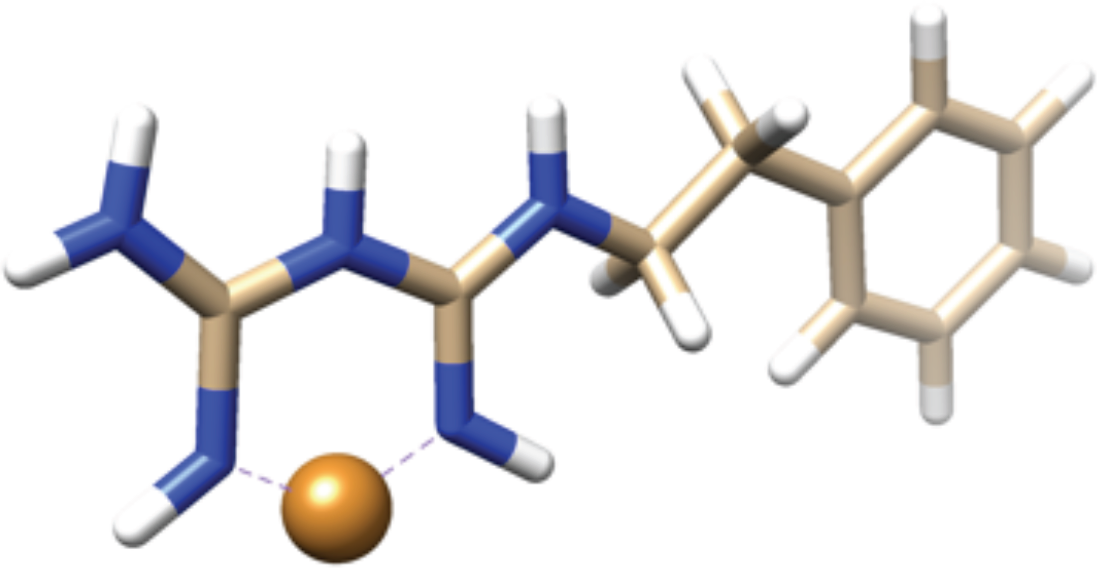
Monophenformin copper complex tridimensional representation.

The complex is imagined to be furtherly coordinated, in solution, with two molecules of water, that are displaced by the cavity residues at the moment of docking.

The results of the docking of the monophenformin copper complex with the cavity’s large entrance, in correspondance to the full slid in FAD pose, are summarised in Table 3.10. No significant interaction was shown with the cavity’s small entrance.

**Table 3.10.** Results of the phenformin molecular docking with the small entrance with different FAD poses.

| FAD position | Phenformin binding energy/kcal mol<sup>-1</sup> | Phenformin inhibition constant(KI)/nM | Hydrogen bonds | Other bonds |
| --- | --- | --- | --- | --- |
| **Fully slid in** | NA – The small cavity is occupied by FAD | NA – The small cavity is occupied by FAD | - | - |
| **Slightly slid out** | -5.7 | 6.60~~×~~ 10<sup>4</sup> | 2 | 3 |
| **Significantly slid out** | -6.0 | 3.97~~×~~ 10<sup>4</sup> | 1 | 7 |
| **Extremely slid out** | -6.3 | 2.39~~×~~ 10<sup>4</sup> | 3 | 7 |

The interactions of the complex with the cavity residues are shown in Figure 3.33.

**Figure 3.33.**
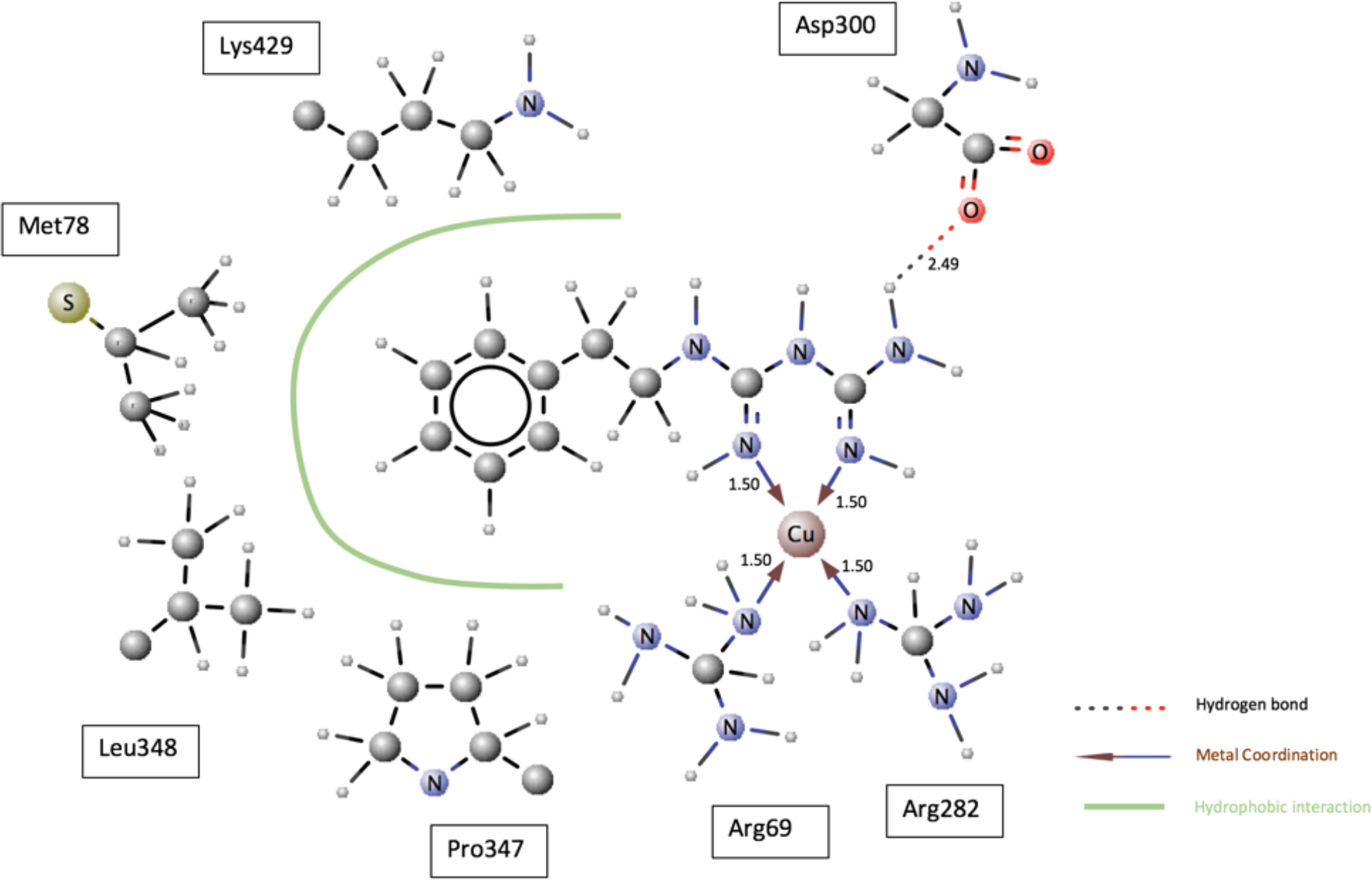
Monophenformin copper complex interactions with cavity residues, bond lengths are expressed in angstroms.

When the pose of the monophenformin copper complex docked with the fully slid in FAD and the pose of the slightly slid out FAD are superimposed a collision is shown, demonstrating that the complex is able to block the FAD sliding out movement. This is shown in Figure 3.34.

**Figure 3.34.**
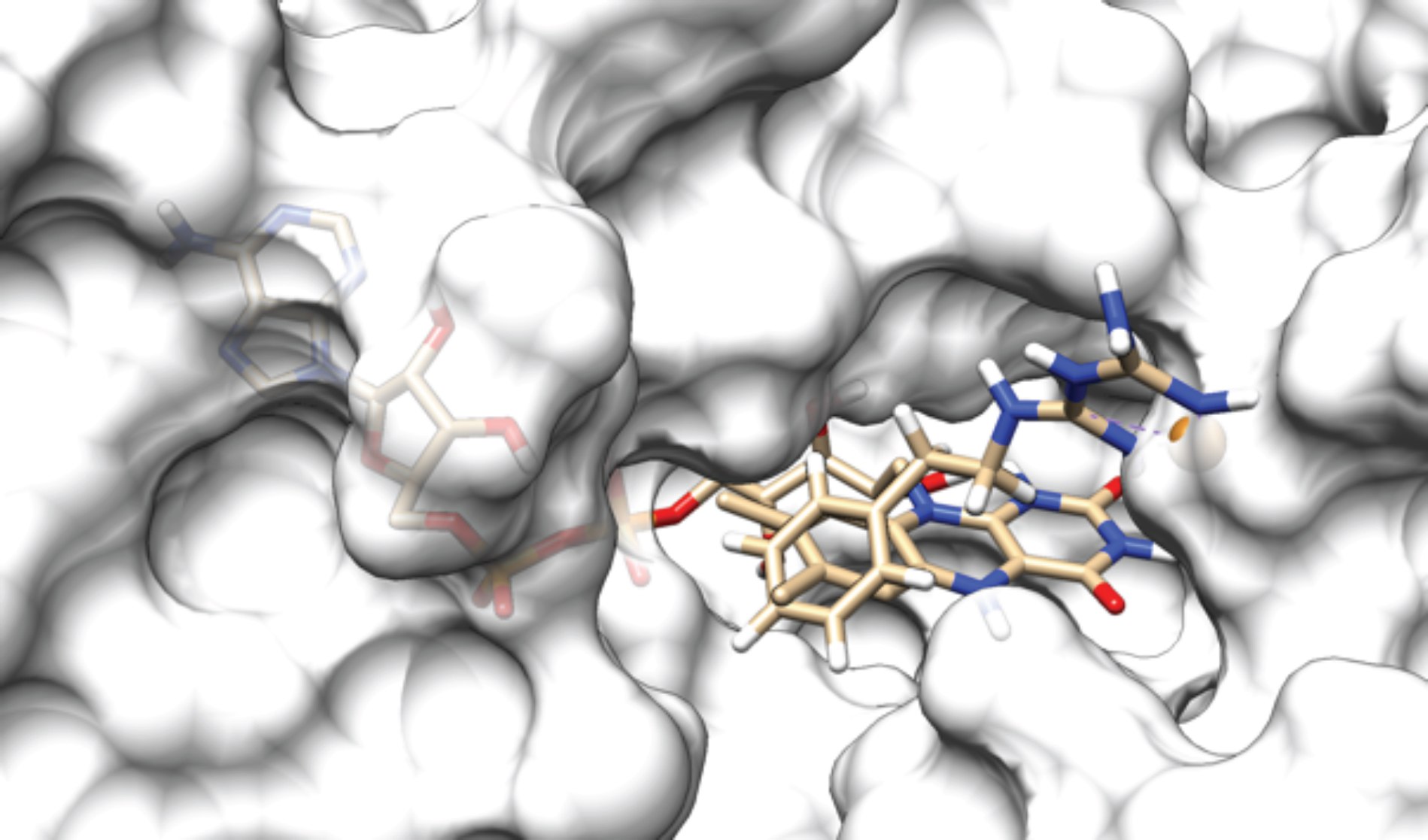
Monophenformin complex collision with slightly slid out FAD.

The more protruding FAD positions are blocked as well.

#### 3.2.10 Monometformin copper complex at the large entrance

In analogy with the monophenformin simulation, a mono-deprotonated monometformin copper complex (Figure 3.35) was docked to both the small and the large entrance of the FAD cavity.

**Figure 3.35.**
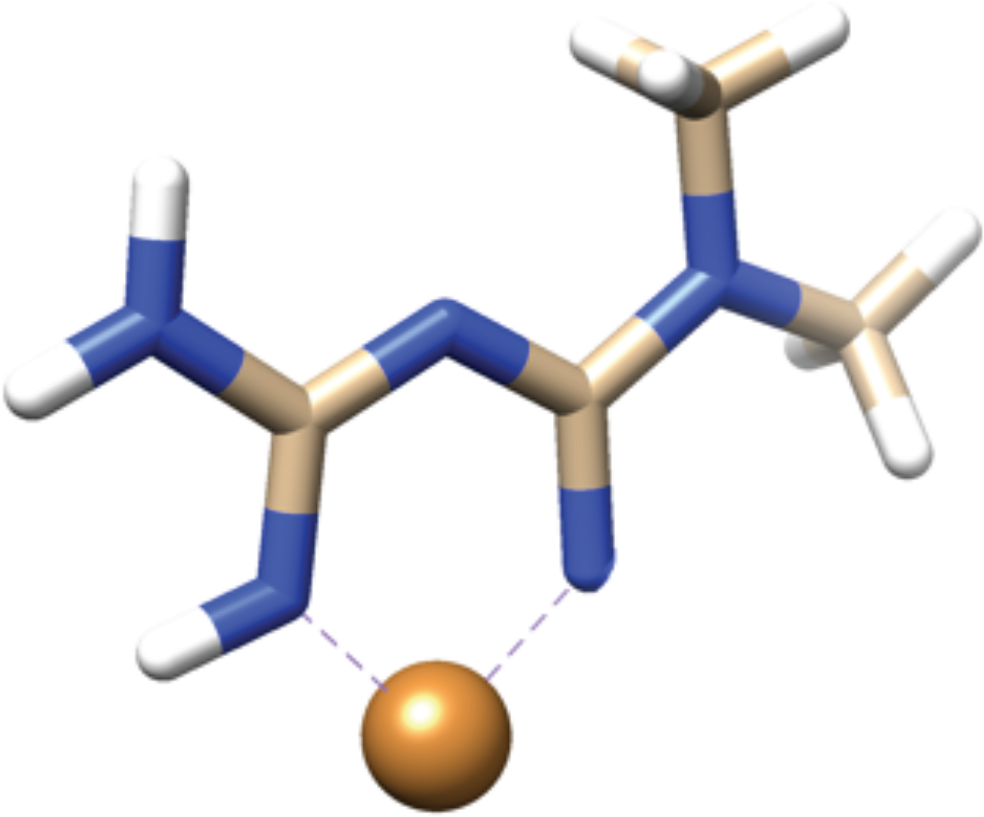
Monometformin copper complex tridimensional representation.

The complex is imagined to be furtherly coordinated, in solution, with two molecules of water, that are displaced by the cavity residues at the moment of docking.

The results of the docking of the monometformin copper complex with the cavity’s large entrance, in correspondance to the full slid in FAD pose, are summarised in Table 3.11. No significant interaction was shown with the cavity’s small entrance.

**Table 3.11.** Results of the monophenformin copper complex molecular docking with the large entrance with fully slid in FAD.

| Copper complex | Binding energy/kcal mol<sup>-1</sup> | Inhibition constant(KI)/nM | Hydrogen bonds | Coordination bonds | Hydrophobic interactions | Ability to prevent docking of Q10 in the cavity |
| --- | --- | --- | --- | --- | --- | --- |
| **[Cu(Phen)]<sup>2+</sup>** | -5.8 | 4.71~~×~~ 10<sup>4</sup> | 1 | 2 | 4 | yes |

The interactions of the complex with the cavity residues is shown in Figure 3.36.

**Figure 3.36.**
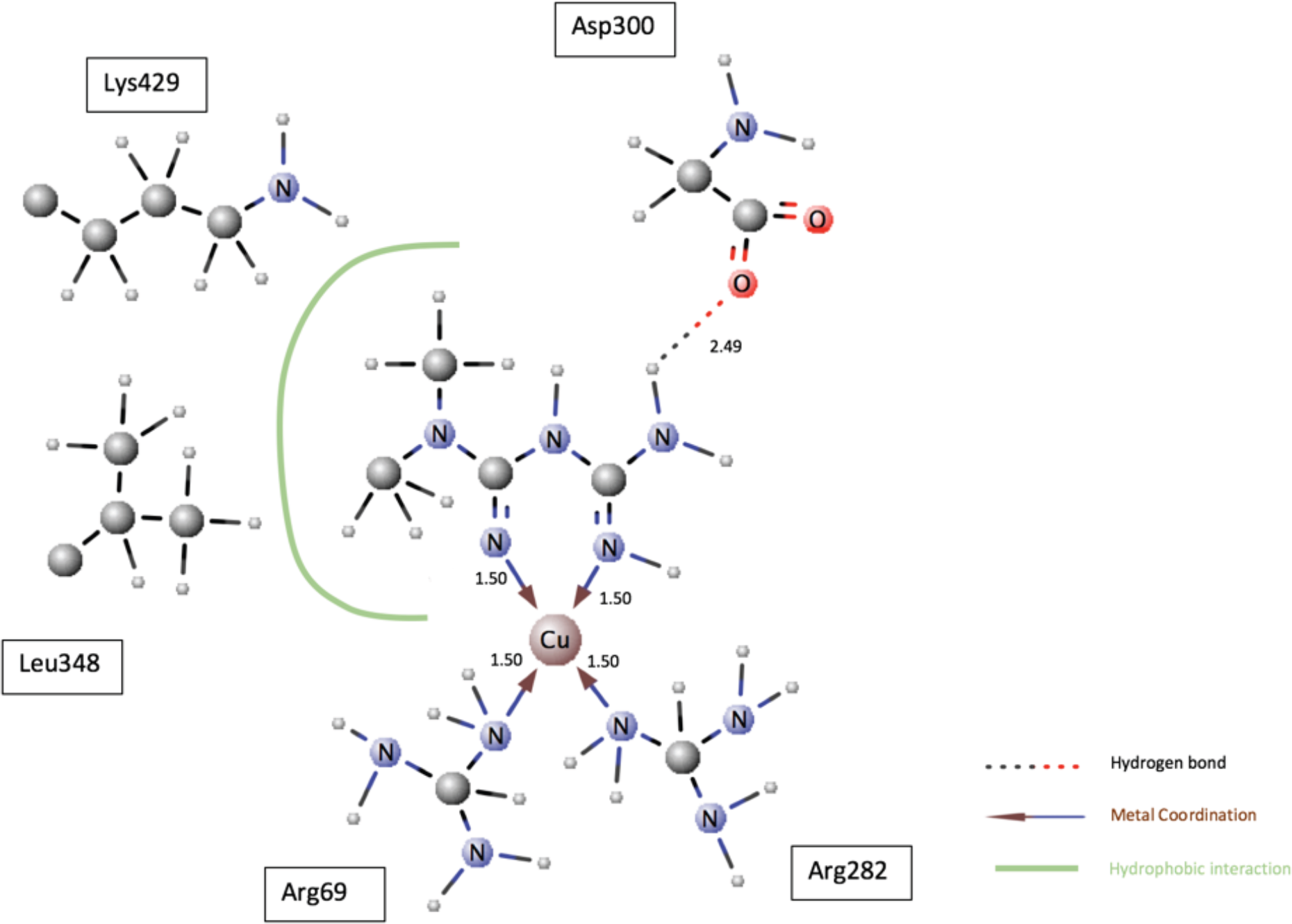
Monometformin copper complex interactions with cavity residues, bond lengths are expressed in angstroms.

When the pose of the monometformin copper complex docked with the fully slid in FAD and the pose of the slightly slid out FAD are superimposed a collision is shown, demonstrating that the complex is able to block the FAD sliding out movement. This is shown in Figure 3.37.

**Figure 3.37.**
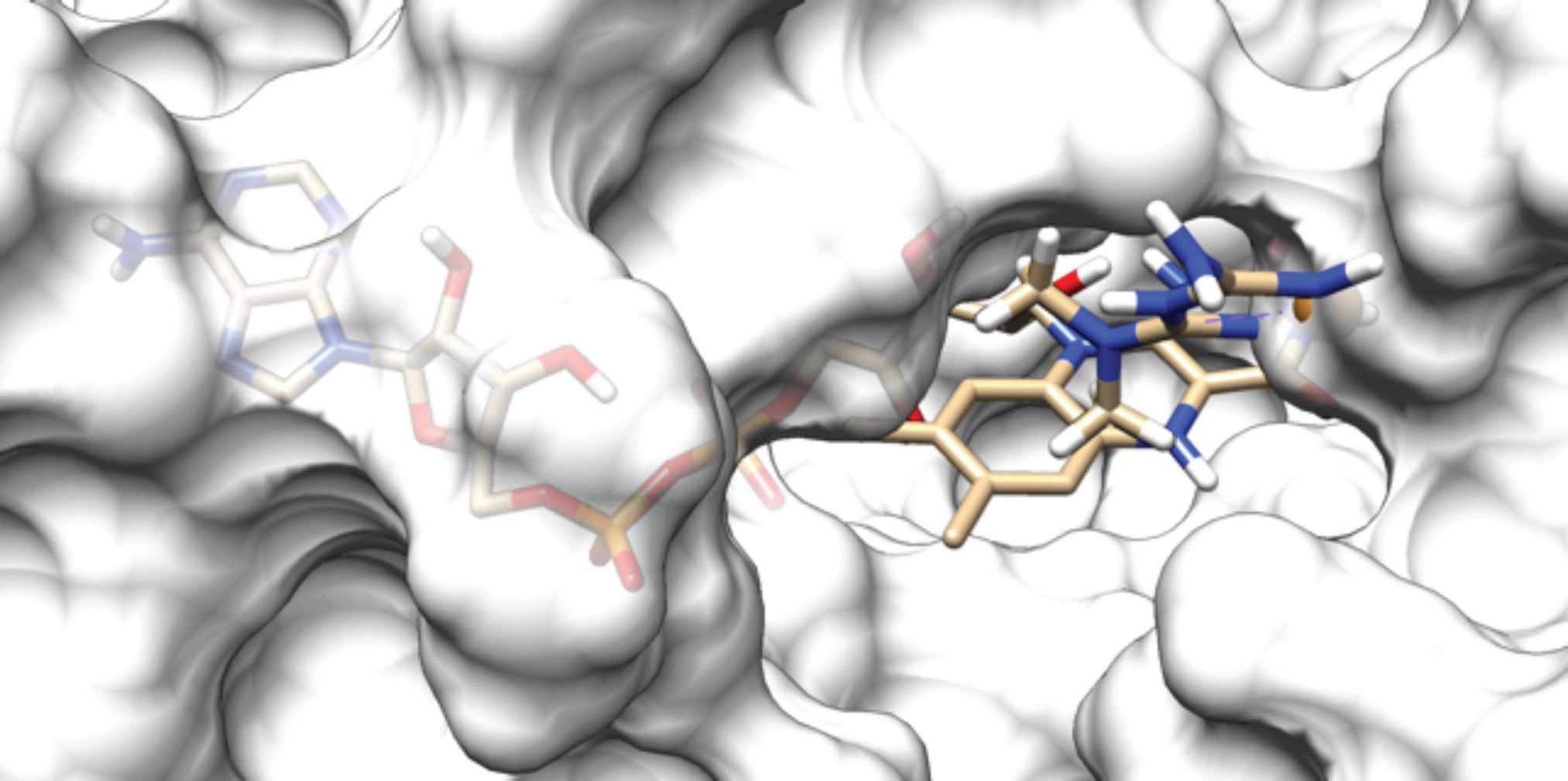
Monometformin complex collision with slightly slid out FAD.

The more protruding FAD positions are blocked as well.

A final simulation showed the same non-interference with G3P docking, as already observed for [Cu(Met-H)_2_]. In Figure 3.38 G3P can be seen docked when [Cu(Met-H)]^1+^ is present.

**Figure 3.38.**
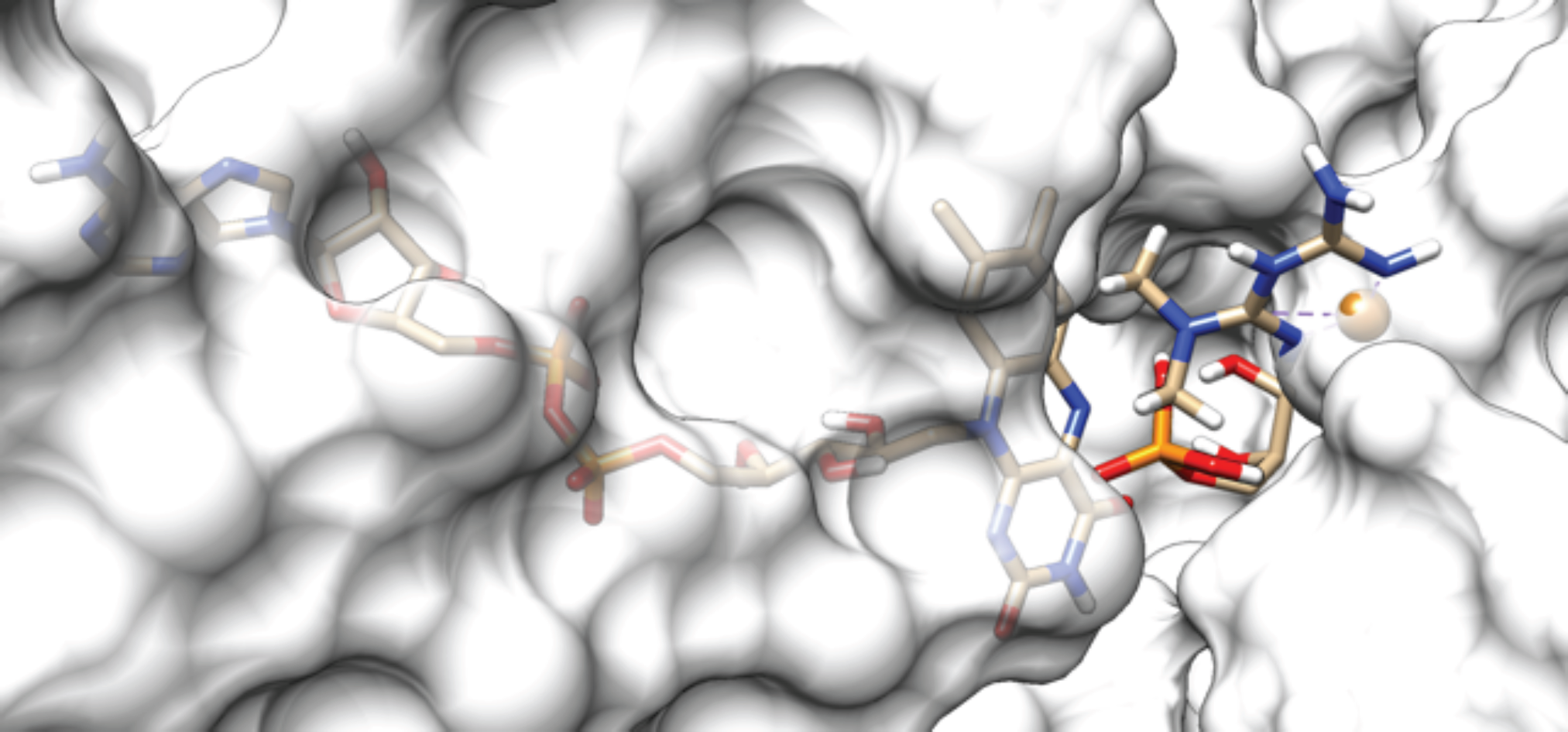
G3P docked to FAD in fully slid in position when [Cu(Met-H)]^1+^ is present.

### 3.3 Virtual screening

#### 3.3.1 Leads identification

The pharmacophore modeled as a result of the analysis of the binding cavity interactions of [Cu(Metf- H)_2_] is represented in Figure 3.39, where the colors used are the standards for H-bond donor, H-bond acceptor, hydrophopic domain and aromatic ring functions.

**Figure 3.39.**
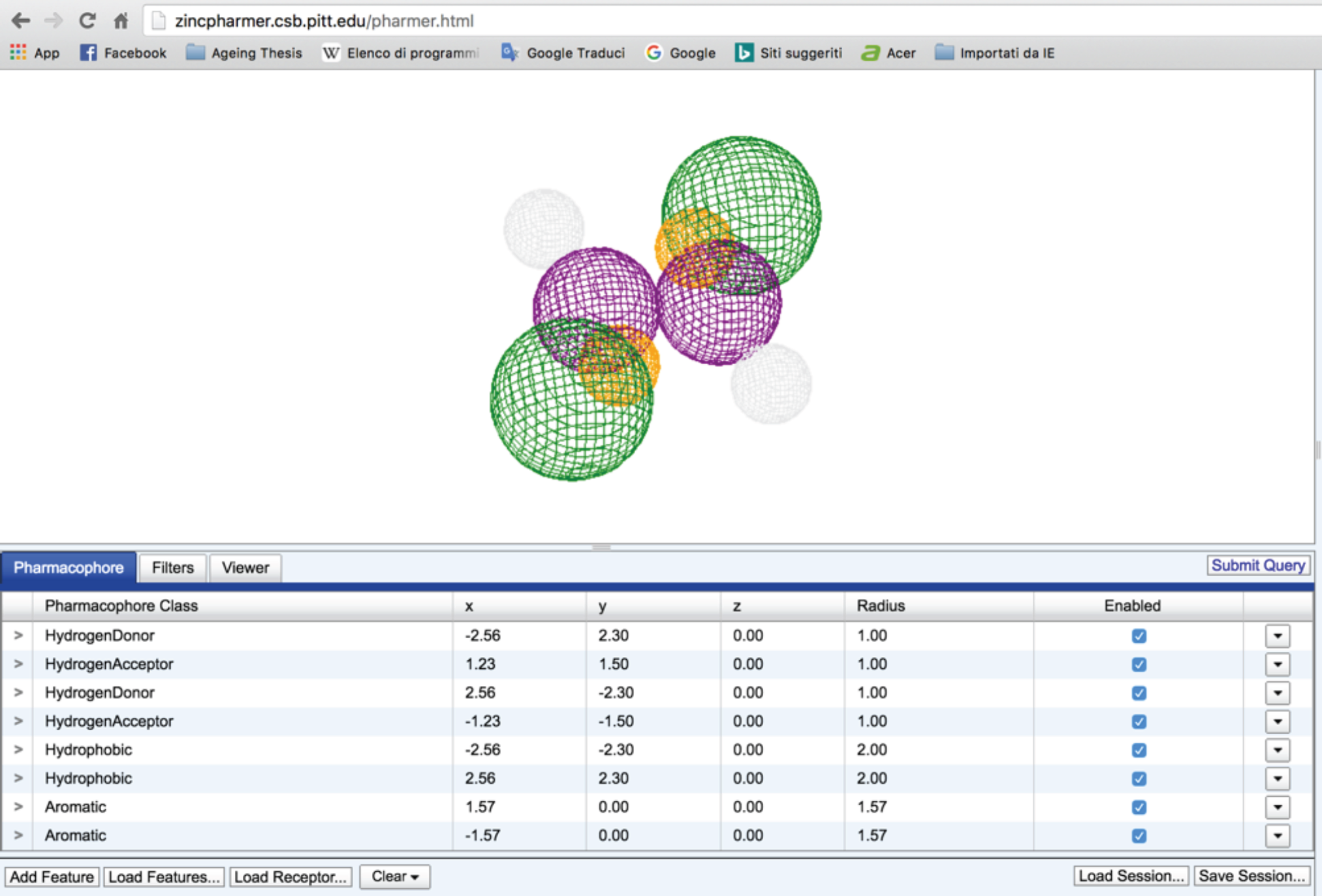
[Cu(Metf-H)_2_] pharmacophore shape and data as modeled on Zinc Pharmer.

The first virtual screening query returned 100 results, the only one fullfilling all the pharmacophore requirements was [4-amino-6-(isopropylthio)-2-methyl-pteridin-7-yl]amine, shown in Figure 3.40. A second search was performed too, this time on ChemSpider (Royal Society of Chemistry, 2016) based on similarity considerations with the monometformin and the monophenformin complexes.

**Figure 3.40.**
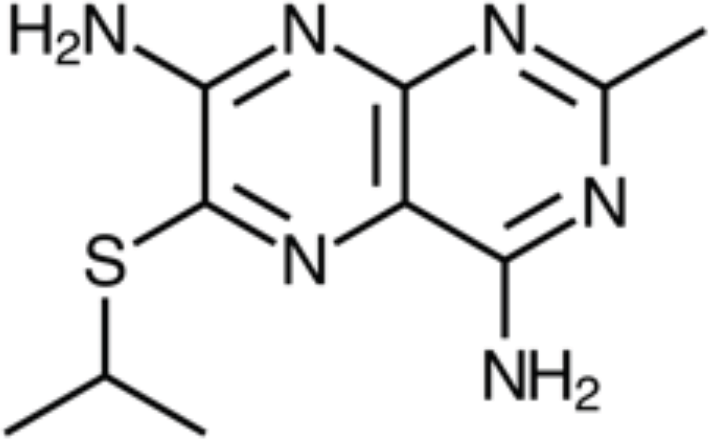
Structure of [4-amino-6-(isopropylthio)-2-methyl-pteridin-7-yl]amine.

In more detail, the compound sought after had the following characteristics:

1. the ability to coordinate in a square planar fashion with Cu^2+^ or Ni^2+^, possibly with stronger ligands than the amine groups, after consulting the NIST database (Smith & Marshall, 2004) the choice fell upon a dithioic acid functional group
2. the ability to perform a hydrogen bond donor function close to the coordinating group
3. the presence of a hydrophobic tail at least four carbons long

The search result was butyl carbamodithioic acid, represented in Figure 3.41.

**Figure 3.41.**
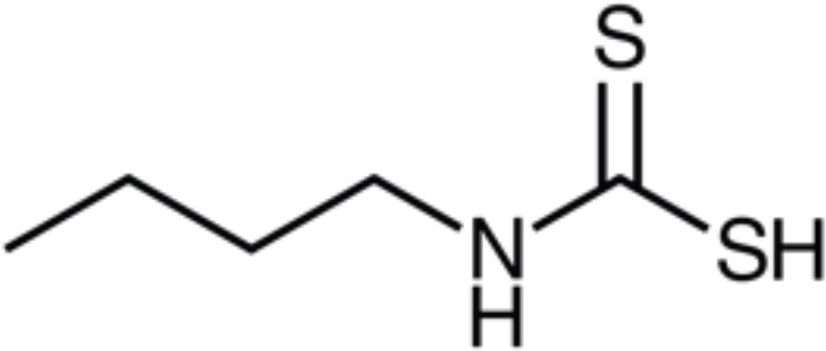
Structure of butyl carbamodithioic acid.

#### 3.3.2 Docking of the virtual screening leads

The results of the docking of the two leads to the cavity’s large entrance are summarised in Table 3.12.

**Table 3.12.** Results of the monometformin copper complex molecular docking with the large entrance with fully slid in FAD.

| Copper complex | Binding energy/kcal mol<sup>-1</sup> | Inhibition constant(KI)/nM | Hydrogen bonds | Coordination bonds | Hydrophobic interactions | Ability to prevent docking of Q10 in the cavity |
| --- | --- | --- | --- | --- | --- | --- |
| **[Cu(Met-H)]<sup>1+</sup>** | -5.7 | 6.60×10<sup>4</sup> | 1 | 2 | 2 | yes |

[4-amino-6-(isopropylthio)-2-methyl-pteridin-7-yl]amine, thanks to its high number (4) of hydrogen bond acceptors, formed a netword of bonding that was even better than the one formed by the original copper complex, with a final binding energy result slightly worse than [Cu(Metf-H)_2_] probably because of the torsional energies involved in the adjustment of the cavity residues around the ligand. [Ni(BCA-H)]^1+^, instead, was slightly penalised with respect to the monometformin copper complex probably because its slimmer shape caused the interaction with some residues to be weaker, for instance the hydrogen bond formed with ASP300 resulted 0.1 angstroms longer (2.6 against 2.5). The detailed interaction diagram of the ligands [4-amino-6-(isopropylthio)-2-methyl-pteridin-7- yl]amine and of [Ni(BCA-H)]^1+^ with each residue are shown in Figures 3.42-3.43.

**Figure 3.42.**
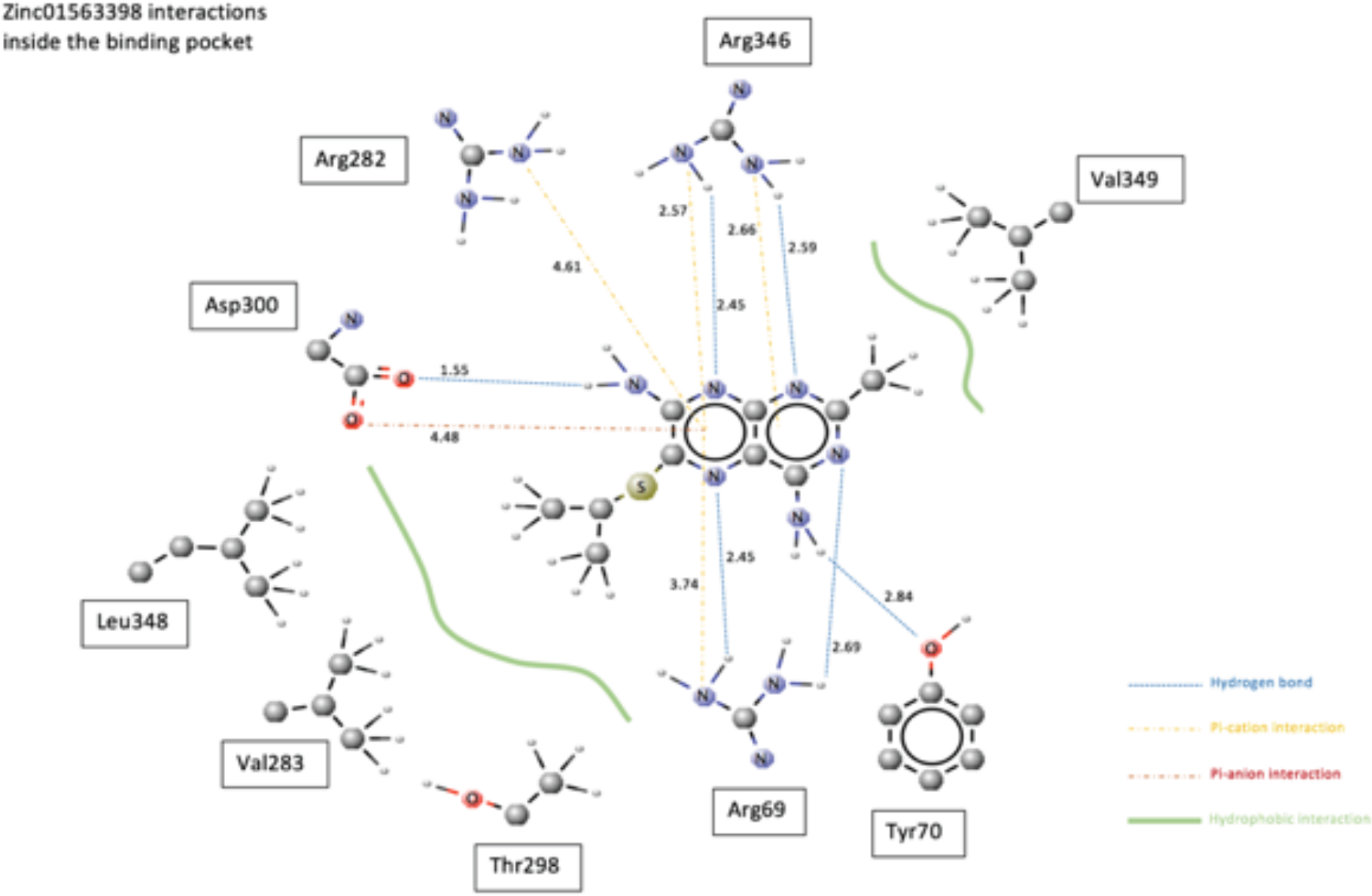
[4-amino-6-(isopropylthio)-2-methyl-pteridin-7-yl]amine’s 2D interaction diagram, bond lengths are in angstroms.

**Figure 3.43.**
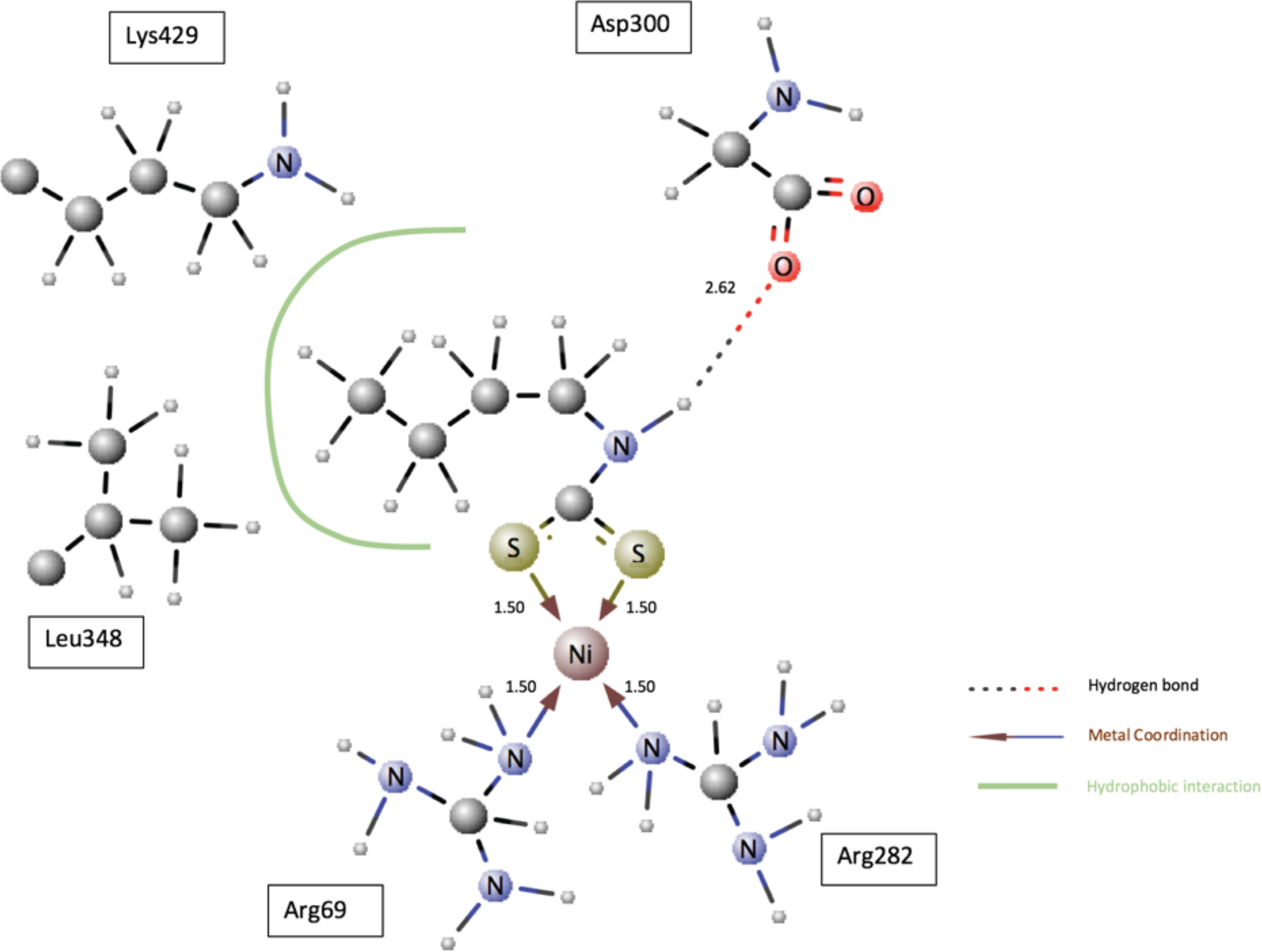
butyl carbamodithioic acid’s 2D interaction diagram, bond lengths are in angstroms.

## Section 4 Discussion

The 2RGH cavity presents a good similarity with the GPD2 sequence with 64% residues in common (Figure 3.1), and the docking of FAD in the modeled cavity shows a RMSD between the original and the new FAD positions of just 0.2 angstroms (Figure 3.4). The modeled cavity can therefore be considered a good approximation of the real one.

FAD’s fully slid in position must be the environment where G3P’s oxidation takes place, because of the following facts:

1. it corresponds to FAD’s lowest binding energy to GPD2 (Table 3.3), therefore is the stablest and the most suited to accept a substrate;
2. it corresponds to G3P’s lowest binding energy to GPD2 (Table 3.4), therefore it’s where G3P spends most of its time on GPD2;
3. it’s the only position characterised by the presence of a hydrogen bond donor close to position 5 nitrogen (Table 3.5), a characteristic needed to initiate the reaction (Ghanem & Gadda, 2008);
4. In that position G3P is oxidised well below the protein surface, exactly what flavoproteins do to protect reactants from solvent (Fraaije & Mattevi, 2000).

The above facts rule out (Madiraju et al., 2014)’s hypothesis that metformin can exert its action at the cavity’s small entrance, because metformin doesn’t bind to the small entrance when FAD is in the fully slid in position (Table 3.7), and this is exactly the moment when G3P binds best (1.-4. above): but a non-competitive inhibitor must bind the receptor equally well both when the substrate is and is not docked, and not only when the substrate (G3P) is away (Strelow et al., 2012).

Both [Cu(Met-H)]^1+^ and [Cu(Met-H)_2_], instead, through binding at the cavity’s large entrance show the ability to interfere with the post G3P oxidation steps, and therefore with the subsequent electron transfer, in one or both of the following ways:

I. preventing the sliding out movement of FADH^-^, needed to reach the slightly slid out position where the negative charge is stabilised by HIS259 (Table 3.5; Fraaije & Mattevi, 2000; Ghanem & Gadda, 2008), a step needed before a proton is taken from the environment and complete reduction to FADH_2_ is finalised and subsequent electron transfer takes place;
II. preventing coenzyme Q10 docking in front of FADH_2_ and subsequent FADH_2_’s oxidation and electron transfer (Tables 3.8 and 3.11).

Furthermore, both complexes have inhibition constants inside (Table 3.8) or close (Table 3.11) to (Madiraju et al., 2014)’s Ki measured range of (3.8 – 5.5) × 10^4^*nM*, they dock effectively only when in deprotonated forms (Tables 3.9 & 3.12) so at high pH values, and they don’t interfere with G3P binding (Figures 3.28 & 3.38), satisfying the requisites of a non-competitive inhibitor.

**Table 3.13.**
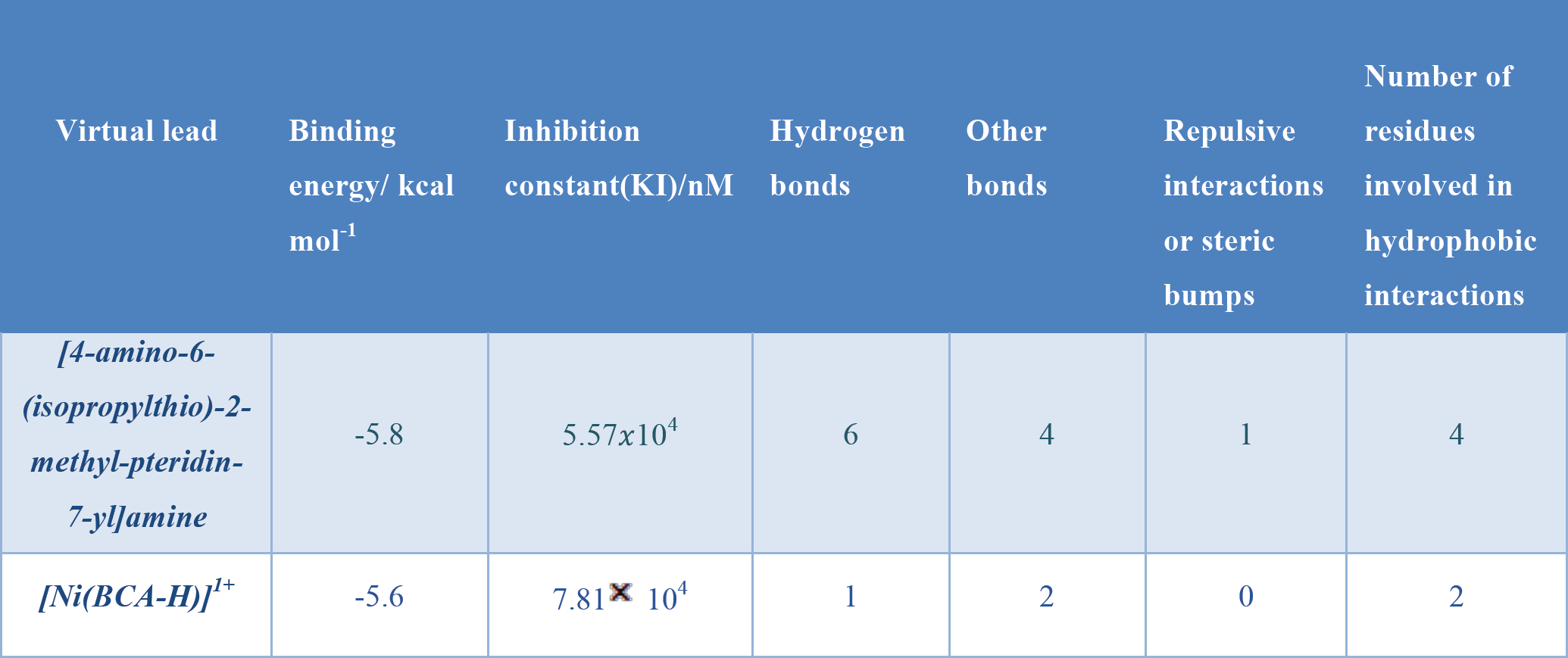
Results of docking of the virtual screening leads in the GPD2 cavity with FAD in fully slid in position.

So in both cases I. and II. above, the suggested mechanisms are able to fit with both with the experimental data of (Madiraju et al., 2014) and with the ideas that mitochondrial high pH (Repiščák et al., 2014), copper (Logie et al., 2012; Annapurna & Rao, 2007) and pseudo-aromaticity in metformin complexes (Repiščák et al., 2014), must be involved.

The model is able to explain in a similar way (Table 3.10 & Figure 3.31) also the antihyperglycemic action of phenformin, a fellow biguanide and predecessor of metformin as an antidiabetic.

In conclusion, this computational study suggests that metformin performs its inhibitory action on gluconeogenesis by sequestering endogenous cellular copper and docking as a metal complex at mitochondrial glycerophosphate dehydrogenase in correspondence of the FAD cavity’s large entrance, this way blocking the electron transfer mechanism taking place after G3P oxidation.

The proposed alternative drugs resulting from the virtual screening represents a possibility to shed light on the issue directly in the laboratory, since evidence of an inhibitory action of [4-amino-6- (isopropylthio)-2-methyl-pteridin-7-yl]amine would confirm the mechanism suggested for the dimetformin complex, while an inhibitory action of [Ni(BCA-H)]^1+^ would confirm the mechanism suggested for the monometformin complex.

Needless to say, a better understanding of metformin mechanism of action would open the possibility to design new antidiabetic drugs taylormade to the characteristics of the target cavity.

## List of abbreviations

ACC = Acetyl-CoA carboxylase;

ADP = Adenosine diphosphate;

AMP = Adenosine monophosphate;

AMPK = 5’ adenosine monophosphate-activated protein kinase;

ATP = Adenosine triphosphate;

BCA = Butyl carbamodithioic acid;

cAMP = Cyclic adenosine monophosphate;

FAD = Flavin adenine dinucleotide;

FASTA = FAST-All, is a text-based format for representing either nucleotide sequences or peptide sequences (“All” means that it works with any alphabet), in which nucleotides or amino acids are represented using single-letter codes;

G3P = Glyceraldehyde 3-phosphate;

GPD2 = Glycerophosphate dehydrogenase 2, the mitochondrial glycerophosphate dehydrogenase;

KI = Inhibition constant;

LKB1 = Liver Kinase B1;

LOMETS = Local Meta-Threading-Server, is an on-line web service for protein structure prediction;

Metf = Metformin;

Metf-H = Deprotonated metformin, where the deprotonation occurs at the secondary amine in position 2 of the parent chain;

MOL2 = Molecular file format 2;

NIST = National Institute of Standards and Technology;

PDB = Protein Data Bank file format;

PDBQT = Protein Data Bank, Partial Charge (Q), & Atom Type (T)) format; Phen = Phenformin;

Phen-H = Deprotonated phenformin, where the deprotonation occurs at the secondary amine in position 2 of the parent chain.

SDF = Structure data format file;

SREBP-1 = Steroid regulatory element binding protein 1.

